# Correlated Evolution of two Copulatory Organs via a Single Cis-Regulatory Nucleotide Change

**DOI:** 10.1101/313197

**Authors:** Olga Nagy, Isabelle Nuez, Rosina Savisaar, Alexandre E. Peluffo, Amir Yassin, Michael Lang, David L. Stern, Daniel R. Matute, Jean R. David, Virginie Courtier-Orgogozo

**Author notes:** Correspondence, @Biol4Ever. current address: The Milner Centre for Evolution, Department of Biology and Biochemistry, University of Bath, Bath BA2 7AY, UK.

## Abstract

**One Sentence Summary:** We identify one nucleotide substitution in a gene regulatory region contributing to evolutionary change of two distinct copulatory organs.

**Highlights:**

- We identify a gene and 3 substitutions causing genital evolution between species
- The evolved mutations lie in a pleiotropic enhancer
- One mutation decreases genital bristle number and increases leg sex comb tooth number
- This mutation disrupts a binding site for Abd-B in genitals and for another factor in legs

**SUMMARY:** Diverse traits often covary between species. The possibility that a single mutation could contribute to the evolution of several characters between species is rarely investigated as relatively few cases are dissected at the nucleotide level. *Drosophila santomea* has evolved additional sex comb sensory teeth on its legs and has lost two sensory bristles on its genitalia. We found that a single nucleotide substitution in an enhancer of the *scute* gene contributes to both changes. The mutation alters a binding site for the Hox protein Abdominal-B in the developing genitalia, leading to bristle loss, and for another factor in the developing leg, leading to bristle gain. Our study shows that morphological evolution between species can occur through a single nucleotide change affecting several sexually dimorphic traits.

## RESULTS AND DISCUSSION

### “Variability is governed by many unknown laws, of which correlated growth is probably the most important”[1]

Correlated evolution of traits is widespread among taxa [1,2] and can be due to pleiotropy, where a single locus causally affects several traits [3]. Pleiotropy imposes large constrains on the paths of evolution [4,5], making it crucial to assess the extent of pleiotropy to understand the evolutionary process. Empirical studies suggest that many loci influence multiple traits [3,6,7] and current data cannot reject the idea that all genetic elements have pleiotropic roles [3,8,9]. Several pleiotropic substitutions have been associated with natural variation [10–12, 12b]: most are coding changes and all underlie intraspecific changes (www.gephebase.org). Nevertheless it remains unclear whether pleiotropic mutations contribute also to interspecific evolution, as experimental evidence suggests that the mutations responsible for interspecies evolution may be less pleiotropic than the mutations underlying intraspecific variation [13].

Here we focused on male sexual bristle evolution between *Drosophila yakuba* and *Drosophila santomea*, which diverged approximately 0.5-1 million years ago [14] and can produce fertile F1 females in the laboratory [15], facilitating genetic mapping. We found that hypandrial bristles – two prominent mecanosensory bristles located on the ventral part of male genitalia in all *D. melanogaster* subgroup species – are missing in *D. santomea* males (Fig. 1). Examination of many inbred stocks and 10 closely related species revealed that the absence of hypandrial bristles is a derived *D. santomea-* specific trait (Fig. 1, Tables S1-S2). No other genital bristle type was noticeably variable in number between *D. yakuba* and *D. santomea* (Fig. S1).

**Fig. 1.**
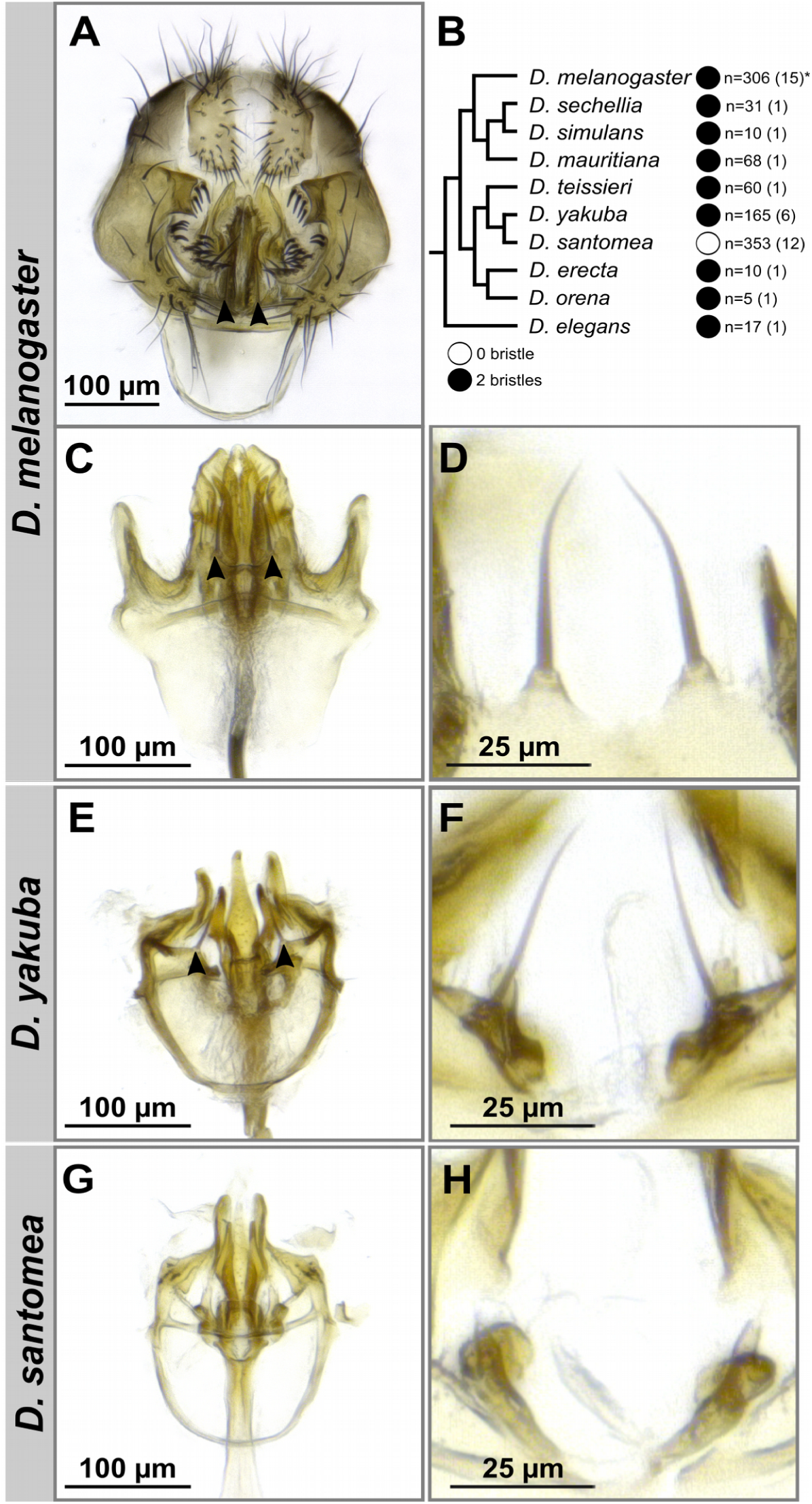
*D. santomea* lost hypandrial bristles. (A) *Drosophila melanogaster* male genitalia. (B) Phylogeny of the *Drosophila melanogaster* species subgroup. All species of have two hypandrial bristles (black circles) except *Drosophila santomea*, which lacks hypandrial bristles (white circle). n: number of scored males, with the number of scored strains in parentheses. Asterisk indicates that 4 males out of 306 had three hypandrial bristles. (C-H) Light microscope preparations of ventral genitalia (C,E,G) and hypandrial bristles (D,F,H) in *D. melanogaster* (C-D), *D. yakuba* (E-F) and *D. santomea* (G-H). Hypandrial bristles are indicated with arrowheads on A, C and E.

We performed whole-genome QTL mapping between *D. santomea* and *D. yakuba* and found that the left tip of chromosome X explains 44% of the variance in hypandrial bristle number in each backcross (confidence interval = 7 Mb for the *D. santomea* backcross and 2.6 Mb for the *D. yakuba* backcross, Fig. 2A). Duplication mapping in rare *D. santomea-D. melanogaster* hybrid males narrowed down the causal region to a 84.6 kb region of the *achaete-scute* complex (*AS-C*) (Fig. 2B-C, Table S2).

**Fig. 2.**
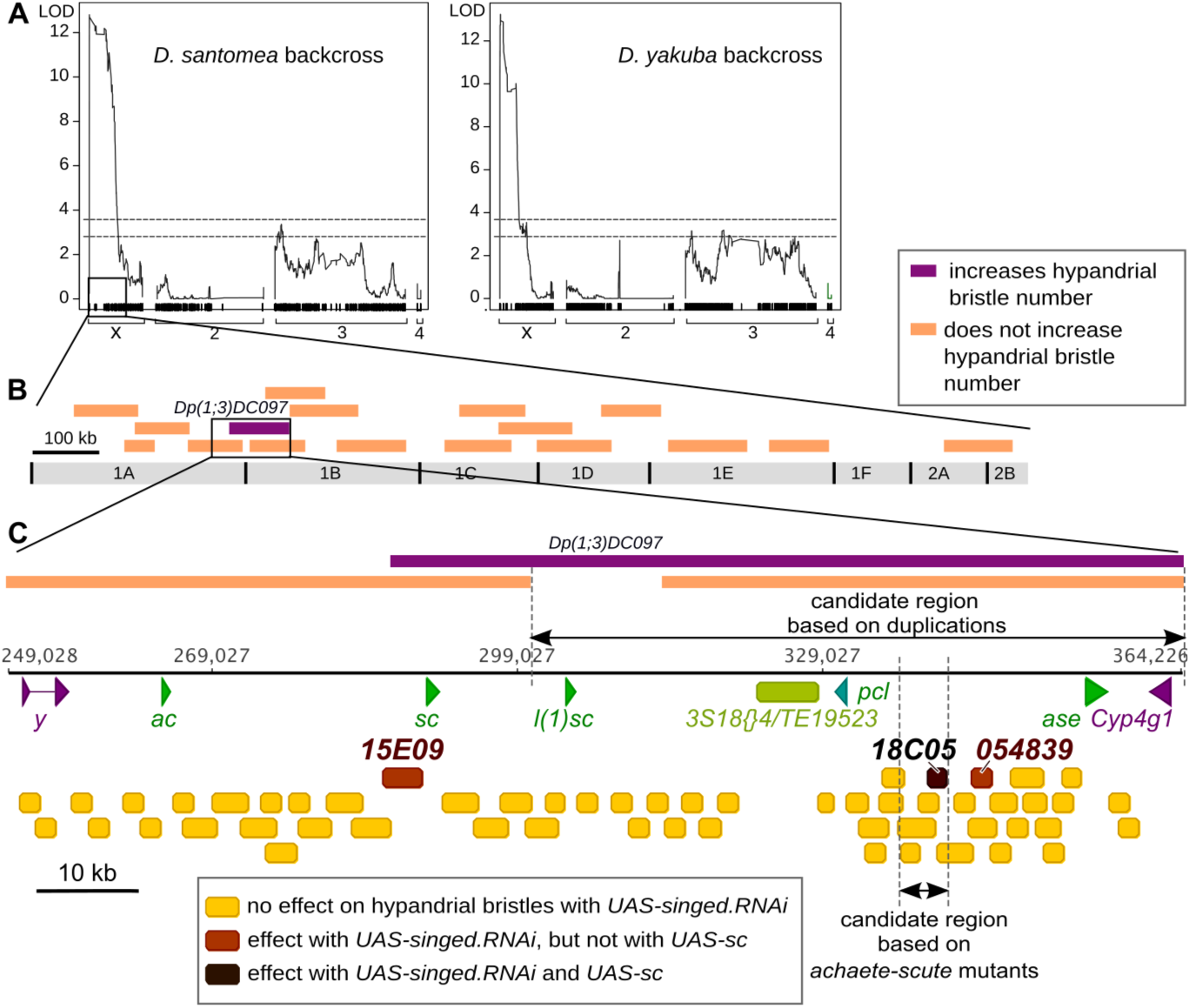
Mapping of the cis-regulatory element involved in hypandrial bristle evolution. (A) QTL analysis of hypandrial bristle number in a *D. santomea* backcross (left) and a *D. yakuba* backcross (right). On the y-axis are the LOD profiles from a Haley-Knott regression analysis. The x-axis represents physical map position in the *D. yakuba* genome. Ticks represent recombination informative markers. Dotted lines represent the 1% (top) and 5% (bottom) significance thresholds. (B) Schematic representation of the left tip of chromosome X and of 19 duplicated fragments of chromosome X that were tested for their effect on hypandrial bristle number in D. santomea-*D. melanogaster* hybrid males. All duplications had no significant effect (orange) except *Dp(1;3)DC097* (purple), which significantly increased hypandrial bristle number. (C) Genomic organization of the *AS*-*C* locus in *D. melanogaster*. Arrows indicate the coding regions of *yellow* (*y*), *achaete* (*a*), *scute* (*sc*), *lethal of scute (l(1)sc), pepsinogen-like (pcl), asense (ase*) and *cytochrome P450-4g1(Cyp4g1*) genes. The light green box represents the insertion of a *3S18{@4/TF9523* natural transposable element. Boxes indicate cis-regulatory elements whose corresponding *GAL4* reporter lines have been tested. Expression of *UAS-singed.RNAi* with 52 *GAL4* lines (yellow boxes) has no effect while it results in singed hypandrium bristles with *15E09*-, *18C05*- and *054839-GAL4*. Extra hypandrial bristles are found with *UAS-sc* and *18C05-GAL4* (dark brown box) but not with *15E09-* and *054839-GAL4* (light brown boxes).

The *AS-C* locus contains four genes, but only two, *achaete (ac*) and *scute* (sc), are required for bristle formation [16]. Both genes are co-expressed, share cis-regulatory elements and act redundantly to specify bristles [17,18]. The elaborate expression pattern of *ac* and *sc* genes prefigures the adult bristle pattern and is controlled by numerous cis-regulatory elements [17]. We tested which of the two genes, *ac* or *sc*, contributes to loss of bristles using null mutants in *D. melanogaster*. All *ac^CAM1^* null mutant males had 2 hypandrial bristles (n=15) and *sc^M6^* null mutants had none (n=15) (Table S4), indicating that *sc* is required for hypandrial bristle development in *D. melanogaster*.

We detected 64 nucleotide differences in the *sc* coding region between *D. yakuba* and *D. santomea*, and all were synonymous substitutions, indicating that coding changes in *sc* are not responsible for the evolved function of *sc*. Using molecularly mapped chromosomal aberrations, we identified a 5-kb region located > 46 kb downstream of the *sc* promoter that is required in *D. melanogaster* for hypandrial bristle development (Fig. S2, Tables S3-4). Independently we screened 55 *GAL4* reporter constructs tiling the entire *AS-C* locus and identified three *GAL4* lines (*15E09, 054839* and *18C05*) that drive expression in hypandrial bristles (Fig. 2C, Tables S5-6). Only one of these lines, *18C05*, increased hypandrial bristle number with *UAS-scute* in a *sc* mutant background or in a *sc+* background (Fig. 2C, Fig. S3E-P). The 2036-bp *18C05* region is located within the 5-kb candidate region identified with *ac-sc* structural mutations (Fig. 2C), suggesting that *18C05* is a good candidate region for hypandrial bristle evolution.

To test whether loss of hypandrial bristles in *D. santomea* resulted from changes(s) in the *18C05 cis*-regulatory region, we assayed whether orthologous *18C05* regions from *D. melanogaster*, *D. yakuba* and *D. santomea* driving a *sc* coding region could rescue hypandrial bristles in a *D. melanogaster sc* mutant (Table S7-8). The *D. melanogaster 18C05* enhancer rescued two bristles in both *sc^29^* and *sc^M6^* mutant backgrounds, indicating that this construct mimics normal levels of *sc* expression. The *D. yakuba 18C05* enhancer rescued on average 2 hypandrial bristles in *sc^M6^* and 0.5 bristles in *sc^29^* whereas the *D. santomea 18C05* enhancer rescued significantly fewer bristles (1.1 in *sc^M6^* and 0 bristles in *sc^29^*, Fig. 3). For another measure of *18C05* enhancer activity, we compared the ability of enhancer-GAL4 constructs containing the *18C05* region from *D. melanogaster, D. yakuba* or *D. santomea* to induce extra bristles in *sc* mutants using the *UAS-GAL4* system with *UAS-sc*. In this assay the *D. santomea 18C05* region also induced fewer bristles than the corresponding *D. yakuba* region (Fig. 3, GLM-Quasi-Poisson, F(19, 509) = 161.7, p < 10^−5^ for *sc^29^;* F(19, 415) = 125.9, p < 10^−5^ for *sc^M6^*). Together, these results suggest that changes(s) within *18C05* contributed to hypandrial bristle evolution in *D. santomea*.

**Fig. 3.**
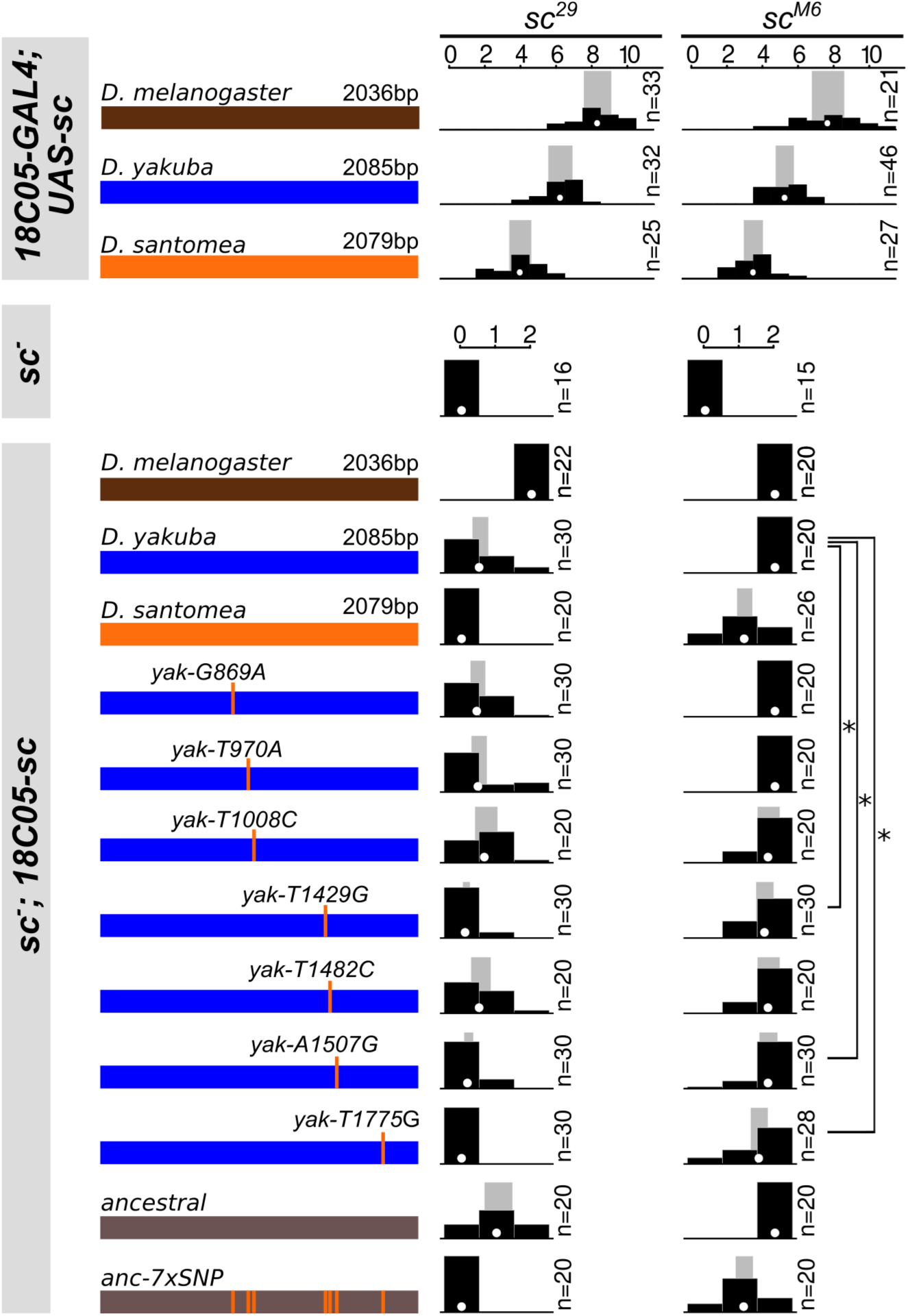
Three *D*. santomea-specific substitutions in *18C05* contribute to the loss of hypandrial bristles. Rescue of the hypandrial bristle loss of *sc^29^* (left column) and *sc^M6^* (right column) *D. melanogaster* mutants by expression of either *GAL4* with *UAS-sc* or *sc* driven by *18C05* sequences from *D. melanogaster* (brown), *D. yakuba* (blue) and *D. santomea* (orange). Seven *D. santomea-specific* substitutions (vertical orange bars) were introduced into either the *D. yakuba* region (blue) or the ancestrally reconstructed *18C05* region (grey). Distribution of hypandrial bristle number (black histogram), together with mean (white dot) and 95% confidence interval (grey rectangle) from a fitted GLM Quasi-Poisson model are shown for each genotype. Note that for a given rescue construct, *18C05-GAL4 UAS-sc* produces more hypandrial bristles than *18C05-sc*, probably due to the amplification of gene expression caused by the *GAL4/UAS* system. n: number of scored individuals. *: p<0.05

To narrow down the region responsible for hypandrial bristle loss, we dissected the *18C05* element from *D. melanogaster, D. yakuba* and *D. santomea* into smaller overlapping pieces and quantified their ability to produce hypandrial bristles with the *GAL4* rescue experiment. For all three species we found that smaller segments rescued significantly fewer bristles than the corresponding full region (Fig. S4-5). Thus, transcription factor binding sites scattered throughout the entire ~2 kb of the *18C05* element are required to drive full expression in the hypandrial bristle region.

Sequence alignment of the *18C05* region from multiple species revealed 11 substitutions and one indel that are fixed and uniquely derived in *D. santomea*. Among them, seven substitutions altered sites that are otherwise conserved in the *D. melanogaster* subgroup (Fig. S6-7). We tested the effect of these seven *D*. santomea-specific nucleotide changes by introducing them one at a time or all together, into either a *D. yakuba 18C05* enhancer or into the inferred ancestral enhancer driving *sc* expression (Tables S7-11). The ancestral *18C05* sequence was resurrected by reverting the *D*. santomea-specific and *D*. yakuba-specific mutations to their ancestral states and it produced the same number of bristles as the *D. yakuba* construct (Fig. 3, Fig. S8). Four substitutions (*G869A, T970A, T1008C* and *T1482C*) had no effect, whether in the *D. yakuba* or in the ancestral background (GLM-Quasi-Poisson, p>0.6). Three substitutions (*T1429G, A1507G* and *T1775G*) decreased the number of rescued bristles in both the *D. yakuba* and the ancestral sequence, and these effects were highly significant, except for *A1507G* in the *D. yakuba* background, which was slightly above statistical threshold (using the most stringent correction method) (Fig. 3). These results are consistent with analysis of smaller pieces of *18C05* and of *18C05* chimeric constructs containing DNA fragments from *D. yakuba* and *D. santomea* (Tables S7-11, Fig. S8). When combined into the *D. yakuba* background, the seven *D*. santomea-specific substitutions rescued the same number of bristles as the *D. santomea 18C05* construct (Fig. 3, Fig. S8, GLM-Quasi-Poisson, p>0.9 in *sc^M6^*). We conclude that at least three fixed substitutions within a 350-bp region located 49 kb away from *sc* contribute to the reduction in hypandrial bristle number in *D. santomea*.

Analysis of *18C05-GAL4* and *18C05-GFP* reporter constructs revealed that the *18C05* region drives expression not only in male genital discs [19] but also in male developing forelegs in the presumptive sex comb region [19b] where *sc* is broadly expressed (Fig. 4A-F). Sex combs are sensory organs used for grasping the female during copulation [20]. They differ in bristle number between *D. santomea* and *D. yakuba* (Fig. S9), and the difference maps to the X chromosome [21], where *sc* is located. These results prompted us to test whether the mutations contributing to hypandrial bristle evolution also affect sex combs. Significantly more GFP-positive cells were detected in the first tarsal segment at 5h after puparium formation (APF) with *18C05yakT1775G-GFP* than with *18C05yak-GFP* (GLM-Poisson, Chi-squared (20,2) deviance = 9.75, p = 0.033), suggesting that *T1775G* increases *sc* expression in the first tarsal segment. Sex comb tooth number was reduced in *sc^M6^* and *sc^6^* mutants and significantly rescued with several *18C05-sc* constructs (Fig. 4J-K). Analysis of *sc^M6^* and *sc^6^* mutants rescued with the *yak18C05-sc* constructs containing the *D. santomea-specific* substitutions showed that *T1429G* and *T1507G* have no effect and that *T1775G* increases the number of sex comb teeth (Fig. 4J-K). We conclude that the *T1775G* substitution contributes to both the increase in sex comb tooth number and the loss of hypandrial bristles.

**Fig. 4.**
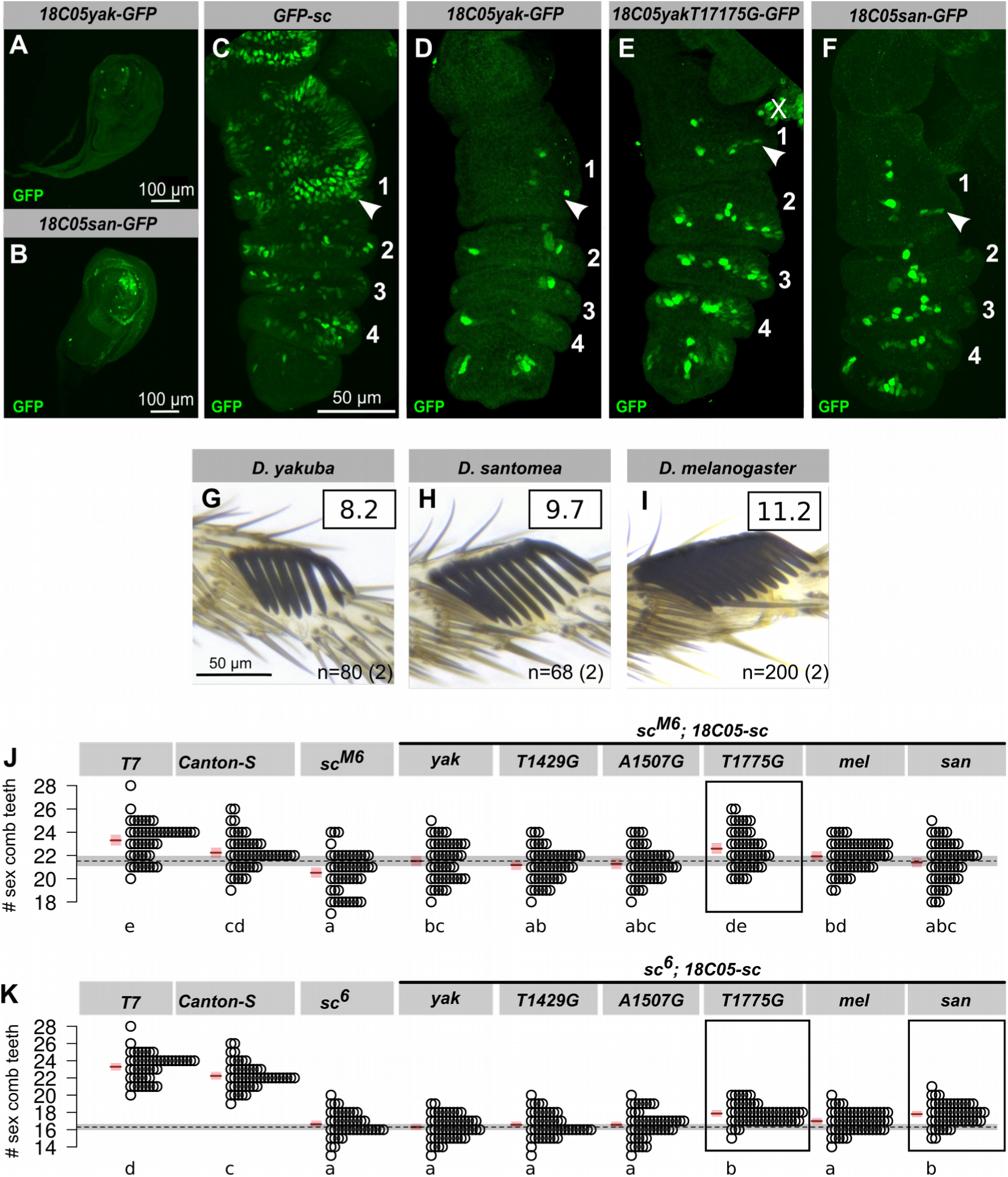
*D. santomea-specific* substitution *T1775G* contributes to increase in sex comb tooth number. (A-F) *18C05* drives expressionGFP staining in T1 leg discs of late L3 larvae (A-B) and in 5h APF pupal legs (C-F) in *D. melanogaster*. Genotype is indicated on top of each panel. Tarsal segments are numbered. Arrowheads point to the presumptive sex comb regions. “X” indicates non-leg tissue. late L3 larvae containing either *18C05yakuba-GFP* (A-C) or *18C05santomea-GFP* (D-F) transgenes. GFP is labeled in green (A, C, D, F), DNA is shown in blue (B, C, E, F). (G-I) Leg sex comb in *D. yakuba* (G), *D. santomea* (H) and *D. melanogaster* (I). Average sex comb tooth numbers per leg are shown in squares. n: number of scored individuals, with the number of scored strains in parentheses. (J-K) Sex comb tooth number in wild-type (T7 and Canton-S), *sc^M6^* (J) and *sc^6^ (K)sc^M6−^* mutants and *sc^M6^* (J) and *sc^6^* (K) and *sc^M6^* mutants rescued with different *18C05-sc* constructs,. Each circle represents one male raised at 25°C. Mean (brown line) and 95% confidence interval (pink rectangle) from a fitted GLM Quasi-Poisson model are shown. Letters indicate the results of all-pairwise comparisons after Holm-Bonferroni correction. Two genotypes are significantly different from each other (p < 0.05) when they do not share a letter. For easier comparison, the horizontal dashed line and the surrounding grey line indicate the mean and 95% confidence interval for *sc^−^;18C05yak-sc*. Transgenic constructs with sex comb tooth number significantly different from 18C05ak-sc are shown in boxes in J-K. On average *D. santomea* males have about 1 extra tooth per sex comb compared to *D. yakuba* (G-H) and 35% of this difference has been attributed to the X chromosome [21]. The substitution *T1775G* produces on average 0.5 extra sex comb tooth per leg, which is more than expected. It is possible that the *D. melanogaster* background, where all our rescue constructs were tested, amplifies the effect of the tested substitutions, especially since *D. melanogaster* males have more sex comb teeth than *D. santomea* or *D. yakuba*.

A bioinformatics search revealed that the *T1775G* substitution is predicted to alter a binding site for the Hox protein Abdominal-B (Abd-B) (Table S12). *Abd-B* is expressed only in the posterior part of the fly, where it directs the development of posterior-specific structures such as the genitalia [22]. We found that reducing *Abd-B* expression, using either genetic mutations or RNA interference, resulted in loss of hypandrial bristles (Fig. S10), indicating that normal levels of *Abd-B* expression are required for hypandrial bristle development. Electrophoretic mobility shift assays showed that Abd-B proteins bind more strongly to a 54-bp fragment of the *18C05* sequence containing the *D*. yakuba-specific T at position 1775 than the *D. santomea-specific* G at this position (Fig. S11). These results are consistent with the hypothesis that the *T1775G* substitution decreases ABD-B binding, contributing to reduction in *sc* expression levels, and ultimately reducing the number of hypandrial bristles. Since *Abd-B* is not expressed in developing legs, *T1775G* is expected to affect binding of other factors to increase sex comb tooth number. Overall, our study suggests that *T1775G* alters overlapping binding sites for distinct factors in the leg and the genitalia (Fig. 4K). All our analyses of the effects of individual substitutions have been carried out in *D. melanogaster* background. It is thus possible that the *18C05* enhancer represents only part of the effect of the *sc* locus on bristle divergence.

Intriguingly, the two organs affected by substitution *T1775G* – hypandrial bristles and sex combs – may both aid the male to position himself on top of the female during copulation [20,23]. Genitals are the most rapidly evolving organs in animals with internal fertilization [24]. To our knowledge, only two other mutations contributing to the evolution of genital anatomy are known. First, a 61-kb-deletion of a cis-regulatory region of the *androgen receptor (AR*) gene in humans is associated with loss of keratinized penile spines in humans compared to chimpanzees [25]. Second, an amino acid change in the *nath10 acetyltransferase* gene which probably appeared recently in laboratory strains of the nematode *C. elegans*, alters morphology in the presence of some mutations but not in a wild-type genetic background [10]. Both mutations appear to be pleiotropic: the *AR* deletion is associated with loss of facial vibrissae in humans and the *nath10* mutation affects egg and sperm production as well. The paucity of known mutations responsible for genital evolution makes it currently difficult to propose general rules for the causes of rapid genital evolution. Our results are reminiscent of Mayr’s pleiotropy hypothesis [26], which posits that certain characters may evolve arbitrarily as a result of selection on other traits due to pleiotropic mutations. In our case, whether the evolutionary change in sex comb tooth number or in genital bristle number has any effect on fitness is unknown.

We report here the first case of a cis-regulatory substitution between species with pleiotropic effects. Given the large number of bristle types regulated by *sc* (>100 in adult flies), it is possible that no cis-regulatory mutation in *sc* can affect only one bristle type. Our results challenge the idea that cis-regulatory enhancers are strict tissue-specific modules underlying evolutionary changes in targeted traits [27]. Even though cis-regulatory mutations may affect several tissues, it is probable that they still tend to be less pleiotropic than coding changes. Our results are thus compatible with the idea that cis-regulatory changes tend to have fewer pleiotropic effects than coding changes on average. Enhancer sequences evolve rapidly, with rapid turn over of individual binding sites while maintaining transcriptional output over millions of years by compensatory mutations [28]. Since pleiotropic mutations can have deleterious off-target effects, we propose that evolution of pleiotropic sites within enhancers should trigger the subsequent selection of compensatory mutations in cis, thus contributing to rapid divergence of cis-regulatory sequences. Overall, our results suggest that pleiotropic cis-regulatory mutations may play a more important role in evolution than previously thought.

## Data and materials availability

Sequences were deposited into GenBank (accession numbers MG460736-MG460765). Source data for Bristle Number and QTL mapping analysis are available as Auxiliary Supplemental Files.

## Supplemental Information includes

Figures S1-S11

Tables S1-S13

Supplemental References (29-63)

## ACKNOWLEDGEMENTS

We thank the Tucson Drosophila Species Stock Center, the VDRC, Kyoto and Bloomington Stock Centers for flies. We thank São Tomé authorities for allowing us to collect flies. We thank S. Picard for help with MSG, E. Sánchez-Herrero for *Abd-B* flies, F. Schweisguth for *GFP-sc* flies, J. Selegue and S.B. Carroll for the *Abd-B* construct, L. Pintard and N. Joly for help in protein purification, R. Mann, S. Feng and G. Rice for EMSA suggestions, F. Mallard and T. Tully for advices on the statistical analyses, J. L. Villanueva-Cañas for help with JASPAR. We thank Q.D. Tran, M. Notin, V. Ludger, A. Matamoro-Vidal, C. Nobre, G. Verebes, S. El Ouisi, A. La, F. Foutel-Rodier, C. Guillard-Sirieix and A. Aydogan for their contributions and C. Desplan, D. Petrov, B. Prud’homme, A. Martin and J.A. Lepesant for comments on the manuscript. We acknowledge the ImagoSeine core facility of the Institut Jacques Monod, member of IBiSA and France-BioImaging (ANR-10-INBS-04) infrastructures. We thank the Courtier-Orgogozo team for providing a stimulating environment and technical support.

## AUTHOR CONTRIBUTIONS

J.R.D. found that *D. santomea* lacks hypandrial bristles and that the trait difference is X-linked, D.L.S. genotyped flies with MSG, A.Y., I.N. and V.C.O. performed the QTL mapping experiment, D.R.M. made the *D. santomea-D. melanogaster* hybrids, I.N. dissected them, O.N. did all other fly crosses and dissected them, O.N., I.N., R.S. and A.E.P. phenotyped >3000 males for hypandrial bristles, O.N. phenotyped all other bristles, O.N. and M.L. did EMSA, O.N. and I.N. constructed the plasmids, O.N. performed immunostainings and microscopy, A.E.P. performed all statistical analyses with feedback from O.N. and M.L., D.R.M. collected wild flies, V.C.O. supervised research, performed bioinformatics sequence analysis and wrote the paper with O.N. All authors provided feedback on the text.

## STAR*METHODS

### CONTACT FOR REAGENT AND RESOURCE SHARING

Further information and requests for resources and reagents should be directed to and will be fulfilled by the Lead Contact, Virginie Courtier-Orgogozo.

### EXPERIMENTAL MODEL AND SUBJECT DETAILS

The origin of all the fly strains used can be found in Tables S1,3,5-6. All flies were cultured on standard cornmeal-agar medium in uncrowded conditions at 25°C unless stated. We used *Canton-S* as a wild-type *D. melanogaster* strain. Transgenic constructs were integrated into the *attP2* landing site in *D. melanogaster w^1118^* by BestGene Inc. Hybrid males between *D. yakuba* and *D. santomea* were obtained by collecting 20 virgin females with 20 males from each stocks and crossing them reciprocally in both directions. At least 10 such crosses were made and flipped every 4-5 days for several weeks. For QTL mapping, *D. yakuba* yellow[1] virgin females were crossed *en masse* to *D. santomea* SYN2005 males to generate F1 hybrid females, which were subsequently backcrossed, separately, to both parental strains. Genitalia of backcross males were isolated for dissection and the remaining carcass was stored at −20 °C for subsequent sequencing library preparation.

### METHOD DETAILS

#### Genotyping of backcross males for QTL mapping

The carcass of each male was crushed in a 1.5-ml Eppendorf tube with a manual pestle in 180 μl of Qiagen Tissue Lysis buffer. DNA of individual flies was extracted using Qiagen DNeasy Blood & Tissue extraction kit (cat #69506). A Multiplexed Shotgun Genotyping sequencing library was made from 189 *D. santomea* backcross males and for 181 *D. yakuba* backcross males as described previously [29]. The list of barcodes used in this study are provided in Supplemental Data File 1, within the names of the individuals that were sequenced. *D. yakuba* and *D. santomea* parental genome sequences were generated by updating the *D. yakuba* genome sequence dyak-4-chromosome-r1.3.fasta with Illumina paired-end reads from *D. yakuba yellow[1]* and *D. santomea* SYN2005 (sequenced by BGI) using the msgUpdateParentals.pl function of the MSG software package. The resulting updated genome files are dsan-all-chromosome-yak1.3-r1.0.fasta.msg.updated.fasta and dyak-4-chromosome-r1.3.fasta.msg.updated.fasta. Ancestry was estimated for all backcross progeny using MSG software (github.com/YourePrettyGood/msg). Ancestry files were reduced to only those markers informative for recombination events using the script pull_thin_tsv.py (github.com/dstern/pull_thin). Markers were considered informative when the conditional probability of being homozygous differed by more than 0.05 from their neighboring markers.

#### QTL mapping

QTL mapping was performed using the R/qtl package version 1.4 [30,31]. The thinned posterior genotype probabilities were imported into R/qtl using the R function read.cross.msg.1.5.R (github.com/dstern/read_cross_msg). QTL mapping was performed independently on each backcross population. We performed genome scans with a single QTL model (“scanone”) using the Haley-Knott regression method [32] performs well with genotype information at a large number of positions along the genome. The genome-wide 5% and 1% significance levels were determined using 1,000 permutations. One QTL peak above the 1% significance level was found for both backcrosses. To check for additional QTL, we built a QTL model with this single QTL using the “fitqtl” function and scanned for additional QTL using the “addqtl” function. A second QTL was found on chromosome 3 for both backcrosses. When introduced into a new multiple QTL model, refined and fitted to account for possible interactions, a third significant QTL was found. Based on the full three-QTL model, no additional significant QTL were found with the function “addqtl”: the highest LOD score for a fourth QTL reached only 1.8 and 1.2 for the *D. yakuba* backcross and the *D. santomea* backcross, respectively. Various three-QTL models with different interactions between loci were assessed. Positive significant interaction was detected between the QTL on chromosome 1 and both QTLs on chromosome 3. The interaction between the two QTLs on chromosome 3 was not significant. For the three-QTL model with interactions between the QTL on chromosome 1 and both QTLs on chromosome 3, we computed the LOD score of the full model and the estimated effects of each locus. The 2-LOD intervals were calculated using the “lodint” function with parameter drop of 2. Analysis of F1 hybrid males is consistent with a large effect of the X chromosome on hypandrial bristle number: male F1 hybrids carrying a *D. yakuba* X chromosome have on average 1.9 hypandrial bristles (n=34) while reciprocal hybrid males possessing the *D. santomea* X chromosome have none (n=29) (Table S2). Note that few informative markers are found on the right arm of chromosome 2, suggesting the presence of an inversion between parental lines. In both backcrosses the large-effect QTL is estimated to cause a decrease of 0.9±0.1 bristles between a *D. yakuba* hemizygote and a *D. santomea* hemizygote male (Data S1). The QTL peak is at position 46,886 and 221,928 for the *D. santomea* and *D. yakuba* backcross, respectively. The *AS-C* locus is at position 179,000-290,000.

#### Duplication Mapping in *D. santomea-D. melanogaster* hybrids

We used a set of *D. melanogaster* duplication lines to test overlapping parts of chromosome X for their effect on hypandrial bristle number [33]. Each line contains a fragment of the chromosome X inserted into the same attP docking site on chromosome 3L using ΦC31 integrase, allowing direct comparison between fragments. Each duplication was used to screen for complementation of the loss of function allele(s) from *D. santomea*. We exploited the fact that rare *D. santomea-D. melanogaster* hybrid males can be produced by crossing *D. melanogaster* females carrying a compound X chromosome with *D. santomea* males [34]. The resulting hybrid males carry a *D. santomea* X chromosome. We first created a *D. melanogaster* stock whose genotype is *TM3, Sb[1] Ser[1]/Nup98-96[339]* by crossing *Nup98-96[339]/TM3, Sb[1]* with *Df(3R)D605/TM3, Sb[1] Ser[1]*. We then performed three successive crosses at room temperature in glass vials: (a) *C(1)RM, y[1] w[1] f[1]; +/+ × +/+; TM3, Sb[1] Ser[1]/Nup98-96[339]*, (b) *C(1)RM, y[1] w[1] f[1]; TM3, Sb[1] Ser[1]/+ × +/Y; Dp(1,3)/Dp(1,3*), (c) *C(1)RM, y[1] w[1] f[1]; TM3, Sb[1] Ser[1]/Dp(1,3) D. melanogaster* females *× D. santomea* males. The same procedure was followed for 21 duplication lines and progeny was obtained for 17 of them. Hybrid males from the last cross were sorted in two pools, the [*Sb^−^*, *Ser^−^*] males who carried the duplication and the [*Sb*^+^, *Ser^+^*] males which were used as controls which carried no duplication but the balancer chromosome *TM3 Sb[1] Ser[1]*. In *D. melanogaster/D. santomea* hybrids, dominant markers are not always fully penetrant. A few progeny males exhibited [*Sb*^+^, *Ser^−^*] or [*Sb*^−^, *Ser^+^*] phenotypes; they were considered as control individuals carrying the balancer chromosome *TM3, Sb[1] Ser[1]*. Males were stored in ethanol until dissection. Duplication mapping narrowed down the causal region to a 84.6 kb region (*DC097*) of the *achaete-scute* complex (AS-C) (Fig. 2.B-C, Table S2, GLM-Poisson, Chisq(17,478)=398.44, p = 10^−4^).

#### Examination of Hypandrial Bristle Phenotypes

Male genitalia were cut with forceps and then hypandria were dissected with fine needles or forceps Dumont #5 (112525-20, Phymep) in a drop of 1x PBS. For *D. melanogaster* in order to see the hypandrial bristles better we removed the aedeagus by holding the aedeagal apodem with forceps and gently pushing the hypandrium upwards with an other forceps until it separated. Hypandria were mounted in DMHF (Dimethyl Hydantoin Formaldehyde, Entomopraxis). Before dissection, males were sometimes stored at −20°C in empty Eppendorf tubes or in glycerol:acetate:ethanol (1:1:3) solution. For analysis of non-hypandrial bristles, males were stored at −20°C in glycerol:acetate:ethanol (1:1:3) solution. We never stored these males in empty tubes because we found that such a storage procedure can break and remove external bristles (but, as far as we know, hypandrial bristles were not affected by such a procedure, maybe because hypandrial bristles are relatively internal and protected by the epandrium). Furthermore, we never observed a single socket devoid of shaft on the male hypandrium, indicating that hypandrial bristles cannot be accidentally cut or lost with our experimental protocol. 3D projection images of the preparations were taken at 500X magnification with the Keyence digital microscope VHX 2000 using optical zoom lens VH-Z20R/W.

#### Examination of Other Bristles

Since genitalia are the most rapidly evolving organs in animals with internal fertilization [32], we compared the number of genital bristles between two strains of *D. yakuba* and two strains of *D. santomea*. We found no difference between *D. yakuba* and *D. santomea* in any genital bristles except for anal plate and clasper bristles, where a slightly significant interspecific variation was detected (Fig. S1.). The loss of hypandrial bristles in *D. santomea* is thus the major change in genital bristles between *D. santomea* and *D. yakuba*. Genitalia were dissected in 1X PBS, hypandria were removed and the epandria were mounted in 99% glycerol. Gentle pressure was applied on the cover-slip with forceps to flatten the preparations in order to see all bristles. Pictures were taken at a 500X magnification with a digital microscope VHX 2000 (Keyence) using lens VH-Z20R/W. Bristles were counted on the images. For sex comb preparations, prothoracic legs were dissected at the coxa with forceps Dumont #5 and were mounted in DMHF (Dimethyl Hydantoin Formaldehyde, Entomopraxis). Images of the sex combs were taken at 1000x magnification with the Keyence digital microscope as written above. Sex comb teeth were counted on the images with Image J [35].

#### Analysis of *scute* coding sequence

The *scute* coding sequence (CDS) of *D. melanogaster* iso-1 was retrieved from FlyBase. We blasted the updated genome sequences of *D. yakuba yellow[1]* and *D. santomea* SYN2005 (see above) with *D. melanogaster scute* coding region and retrieved only one locus in each species. The *scute* coding region was then annotated with Geneious and no intron was found, as in *D. melanogaster*.

#### Screening *as-GAL4* lines for expression in the hypandrium

The *as-GAL4* lines were ordered from VDRC and Bloomington Stock Center (Table S6). Two lines were not available (*GMR1509* and *VT054822*) so we created new transgenic lines for these regions, named *GMR15X09-GAL4* and *VT054822b-GAL4* (see below). Because screens are easier on adults than on genital discs, and also because the exact developmental stage and location of hypandrial bristle development are unknown [36], we decided to look *for* GAL4-triggered phenotypes in adult males. As a readout *of* GAL4 expression, we tested vario*us* UAS lines (*UAS-mCD8-GFP*, UAS-*yellow* i*n a* yellow mutant backgrou*nd*, *UAS-sc*.RN*Ai*, *UAS-achaete*.RN*Ai, UAS-forked*.RN*Ai, UAS-singed*.RNAi) (lines are listed in Table S1 and S5) together *with DC-GAL4*, which drives expression in the dorso-central thoracic bristles [37]. To enhance RNAi potency we also used *UAS-Dicer-2* (35). With U*AS-mCD8-GFP* and U*AS-yellow* the change in fluorescence or color was hardly visible. The most penetrant bristle phenotype was obtained with U*AS-Dicer-2 UAS-singed.RNAi1^05747^* at 29 °C (Table S5). Therefore this line was chosen for screening all the as-*GAL4* constructs.

Five *as-GAL4* males of each *as-GAL4* line were crossed to five *Dcr2; UAS-singed^105747^.RNAi/CyO* virgin females. Crosses were kept at 29 °C. The non-curly males (*Dcr2; UAS-singed^105747^.RNAi/+*; *+/as-GAL4*) were collected for dissection and kept at −20 °C. Hypandrium dissection and image acquisition were performed as indicated above. For each *as-GAL4* line at least 5 genitalia were examined (Table S6).

To test whether the *15E09, 18C05* and *054839* enhancer-GAL4 drive expression in the hypandrial bristle region in absence of *sc*, we crossed five *sc^29^; UAS-scute (III*) females with five males of each respective *GAL4* line, as well as five *sc^M6^/FM7; UAS-scute (III*) females with five males of each respective *GAL4* line. Of the three GAL4 lines, only *18C05* could induce hypandrial bristles with *UAS-sc* in a *sc* mutant background. The *18C05-GAL4* line produced approximately 10 bristles, where normally only two develop, which may reflect the amplification of gene expression that is inherent to the *UAS-GAL4* system. These results suggest that only *18C05* drives sufficiently strong expression in the hypandrial region to alter bristle patterning.

#### Cloning of *enhancers* into pBPGUw and pBPSUw

Enhancers were cloned into the *GAL4* reporter vector pBPGUw using the same strategy as in [37,38]. Enhancer sequences were amplified by Phusion^®^ High Fidelity Polymerase (New England Biolabs) in two steps reaction using the primers and templates listed in Table S7, S8 and S9. PCR products and vectors were purified by Nucleospin Gel and PCR Clean-Up Kit (Machery-Nagel). Clones were purified by E.Z.N.A.^®^ Plasmid Mini Kit I (Omega Bio-tek). All *GAL4* constructs were cloned using the Gateway^®^ system (ThermoFisher Scientific). The enhancer fragments were first ligateded into *Kpn*I and *Hind*III restriction enzyme site of the vector pENTR/D-TOPO (Addgene) (Table S8). Recombination into the destination vector pBPGUw was performed using LR clonase II enzyme mix (Invitrogen) and products were transformed into One Shot^®^ TOP10 (Invitrogen) competent cells. Recombinant clones were selected by ampicillin resistance on Amp-LB plates (100 μg/ml)

The pBPSUw vector was constructed by replacing the *GAL4* cassette of pBPGUw by *scute* CDS. The *scute CDS* was amplified from *D. melanogaster iso-1* with Scute-CDS-Rev and Scute-CDS-For primers and ligated into pGEM-T Easy (Promega). The *sc-CDS* insert was cut out using KpnI and *HindIII* and cloned into KpnI and *HindIII* sites in pBPGUw, thus replacing *GAL4*. The vector was named pBPSUw where “S” stands for *scute. 18C05* sequences from *D. melanogaster, D. yakuba and D. santomea* were cloned into pBPSUw and tested in rescuing hypandrial bristles in *sc* mutants as written above. We found that *18C05* from *D. melanogaster* rescued two hypandrial bristles in both *sc^29^* and *sc^M6^* mutants. *D. santomea 18C05* enhancer rescued fewer hypandrial bristles on average than the *D. yakuba 18C05* region (Fig. 3., bristle number for *D. yakuba 18C05* in *sc^29^* is significantly different from 0 (Exact-Poisson, p < 10^−16^) and bristle number for *D. santomea 18C05* in *sc^M6^* is significantly different from 2 (Exact-Poisson, p =0.0008)).

The *18C05* full length sequences were amplified by PCR from *D. melanogaster iso-1*(BL2057), *D. melanogaster T-7, D. yakuba Ivory Coast* and *D. santomea SYN2005* with the primers listed in Table S7. The PCR products were cloned into *pBPSUw* as described above. Three different *D. melanogaster 18C05* sequences were tested with *UAS-sc* in the hypandrium in *sc^29^* and *sc^M6^. GMR-18C05* (BL2057) was obtained from the Janelia Farm collection [38] and *18C05_BL2057* and *18C05_T7* were cloned in this study. Hypandrial bristle number was found to be significantly higher for *GMR-18C05* than for *18C05_BL2057* and *18C05_T7* in both backgrounds (GLM-Quasi-Poisson, F(2, 63) = 16.88, both p < 10^−6^ for *sc^29^;* F(2, 58) = 20.9, p < 10^−10^ and p < 10-5 for *sc^M6^*). The *GMR-18C05* fragment is inserted in the expression vector 3’-5’ compared to the *D. melanogaster* genome sequence. In contrast, the *18C05_BL2057* and *18C05_T7* are cloned 5’-3’. All the *18C05* constructs we made were inserted in the same orientation, 5’-3’. *GMR-18C05* and *18C05_BL2057* are the same sequences (from *D. melanogaster* Bloomington Stock Center Strain #2057), but cloned in opposite directions. *18C05_T7* contains the *18C05* sequence of *D. melanogaster T.7* strain. Comparing bristle number between *GMR-18C05-GAL4* and *18C05_BL2057-GAL4* shows that the orientation of the cis-regulatory region has an effect on bristle number.

The *18C05-chimera-pBPSUw* constructs were cloned using Gibson Assembly [39] by fusing together different lengths of *18C05* sequences from *D. yakuba Ivory Coast* and *D. santomea SYN2005*. The different chimeras are listed in Table S9. Cloning primers were designed using NEBuilder Tools (http://nebuilder.neb.com/). Primer sequences and templates used in PCR are listed in Tables S7 and S9. To assemble the *18C05* fragments in pBPSUw (Table S10), the vector was linearized by *Aat*II and *Fse*I restriction enzymes (New England Biolabs Inc.). After digestion thermosensitive alkaline phosphatase (FastAP, ThermoFisher Scientific) was added to the reaction to prevent self-ligation of the plasmid. PCR products and the linearized plasmid were isolated from 1% agarose gels and spin column purified. Gibson Assembly was performed as in [39], except that the assembly reactions were incubated at 37 °C for 10 minutes and then 3 hours at 50 °C in a PCR machine. 2 μl of assembly mixtures were transformed into NEB^®^ 10-beta (New England Biolabs Inc.) competent cells and ampicillin-resistant colonies were selected on 100 μg/ml Amp-LB plates. The Gibson Assembly Master-mix was prepared according to [39], its components were purchased from Sigma-Aldrich.

The *18C05-yakubaSNP-pBPSUw* constructs were cloned by Gibson Assembly as described above, except for *18C05yakT1008C* and *18C05yakT1482C* sequences, which were synthesized and cloned by GenScript^^®^^ (Table S11). The *18C05-ancestral* sequences were synthesized and cloned by GenScript^^®^^ into pBPSUw *Aat*II and *Fse*I sites, except for the *18C05_AncG869A*, *18C05_AncT670G* and *18C05_Anc-7SNP* sequences, which were cloned by us by Gibson Assembly into pBPSUw *Aat*II and *Fse*I sites using the 18C05_Ancestral_Gibson_forward and 18C05_Ancestral_Gibson_reverse primers (Tables S7 and S10).

All transgenic constructs were integrated into the *attP2* landing site in *D. melanogaster w^1118^* by BestGene Inc. The *T1775G* substitution affects nucleotide position 447,055 in the Dm6 reference assembly.

#### Genomic DNA preparations for sequencing the *18C05* region

Genomic DNA was isolated with Zymo Research Quick-DNA™ Miniprep Plus Kit from 3 males and 3 females from the *D. yakuba*, *D. santomea* and *D. teisseri* lines listed in Table S12. *18C05* sequences were amplified with San-Yak_lines_sequencing-For and San-Yak_lines_sequencing-Rev primers (Table S7) using Phusion^^®^^ High Fidelity Polymerase (New England Biolabs).

#### Sequence Analysis

Geneious software was used for cloning design and DNA sequence analysis. Nucleotide positions are given according to the alignment of *D. yakuba* Ivory Coast *18C05* sequence with *D. santomea* SYN2005 *18C05* sequence. The *18C05ancestral* sequence of *D. yakuba* and *D. santomea* was reconstructed in Geneious based on the *18C05* sequence alignment of *Drosophila* lines listed in Table S12. Manual parsimony reconstruction of all the ancestral nucleotides was unambiguous, except for one position (766, indel polymorphism), where the sequence is absent in the *simulans* complex and in *D. santomea*, while it is present in *D. teissieri* and polymorphic in *D. yakuba*. For this position we chose *D. teissieri* as the ancestral sequence. The *18C05* sequences of *D. melanogaster* subgroup species were retrieved by BLAST from the NCBI website. Transcription Factor (TF) binding sites in *18C05* were predicted using the JASPAR CORE Insecta database (http://jaspar.genereg.net [40]). 25-60 bp sequences of *18C05* were scanned with all JASPAR matrix models with 50-95% Relative Profile Score Thresholds to test for sensitivity and selectivity [40] (Table S12). For TFs which were absent in JASPAR (Scute), we used Fly Factor Survey to analyze their putative binding affinities to the probe. As *sc* cis-regulatory region is known to contain binding sites for Scute itself [41], we looked for Scute binding sites in *18C05, 15E09* and *054839*. Two putative Scute binding sites (consensus motif *CAYCTGY*, Fly Factor Survey [41]) were found in *15E09* and *054839* but not in *18C05*. Given the present results, we cannot exclude the involvement of *15E09* and *054839* in the evolution of hypandrial bristle evolution in *D. santomea*. In this paper, we decided to focus on the *18C05* enhancer, whose effect could be studied in a *sc* mutant background.

#### Statistical Analyses

Since bristle number is a classical type of count data, we performed statistical analysis using generalized linear models (GLM) and generalized linear mixed models (GLMM) where bristle number, the response variable, is assumed to follow a Poisson distribution [42–44]. All statistical analyses were performed using R 3.4 [45]. GLM were fitted with the function glm() ("stats" core package 3.5.0) and GLMM with the function glmer() ("lme4" package 1.1-14 [46] with the parameter "family" taken to be "Poisson". We tested differences in bristle number by comparing two wild-type stocks of *D. yakuba* with two wild-type stocks of *D. santomea*. We tested the difference between species, using a GLMM of the Poisson type (GLMM-Poisson) where the number of bristles was the response variable, species was a fixed effect to test and stock a random effect. For all other analyses, we tested differences in bristle number between genotypes using GLM of the Poisson type (GLM-Poisson) where the response variable was bristle number and genotype, a categorical variable, was the fixed effect. When we noticed important differences between residual deviance and residual degrees of freedom, we also fitted a quasi-likelihood model of the type "quasi-Poisson" (GLM-Quasi-Poisson) which allows for a model of the Poisson type, but where the variance can differ from the mean and is estimated based on a dispersion parameter (see for example [44] p. 225). For each model, in order to retain the model that fitted best to the data, analysis of deviance was performed using the anova.glm() with "test = Chisq" for GLM-Poisson and "test = F" for GLM-Quasi-Poisson. When needed, we performed multiple comparisons using the glht() function and the "Holm" adjustment parameter ("multcomp" package 1.4-7 [47]) which performs multiple comparisons between fitted GLM parameters and yields adjusted p-values corrected according to the Holm-Bonferroni method [48,49] also performed an exact Poisson test (R function "poisson.test") to test sample mean to a reference value assuming a Poisson distribution. Mean and 95% confidence intervals were directly extracted from the fitted GLM and transformed using exp(coef()) and exp(confint.default()).

For EMSA data, response curves were compared between yak probe and san probe using an ANCOVA after natural log transformation. The unlabeled san 450x responses were compared between yak probe and san probe using a one-sided Mann-Whitney U-test.

#### Abd-B homeodomain (Abd-B-HD) purification and EMSA

The Abd-B-HD-pGEX-4T-1 plasmid (kindly provided by Sangyun Jeong) was transformed into BL21 (DE3) chemically competent cells. Protein expression was induced by 0.1 mM IPTG (isopropyl-β-D-thiogalactopyranoside, Sigma Aldrich). Recombinant protein was purified from 500 ml of bacterial culture as described in Frangioni [50] except that proteins were eluted into 50 mM Tris-HCl, pH 8.0, 500 mM NaCl, 10mM reduced glutation (Sigma-Aldrich, G-4251) and 5% glycerol. Concentrations and purity of the protein were determined by SDS-PAGE and Qubit 2.0 Fluorometer (Life Technologies). Protein aliquots of 20 μl were snap-frozen in liquid nitrogen and stored at −80 °C.

The HPLC-purified biotinylated and non-labelled oligonucleotides (Sigma-Aldrich) were used in PCR to obtain 54 bp probes *yak* and *san* (*san=yakT1775G*) from *18C05yak-pBPSUw* and *18C05yakT1775G-pBPSUw* plasmid templates. Oligonucleotides are listed in Table S7. PCR products were column-purified.

We then used electrophoretic mobility shift assay (EMSA) to test whether the purified Abd-B homeodomain (ABD-B-HD) can bind directly to a 54-bp fragment of *18C05* with the T1775G site at position 13 containing either T (*yak* probe) or G (*san* probe). In each binding reaction, 20 fmol of probes were mixed with the purified ABD-B HD ranging from 0-1.25 μg (0 μg, 0.75 μg, 1 μg and 1.25 μg) in binding buffer containing 10mM TRIS pH 7.5, 50 mM Kcl, 0.5 mM DTT, 6.25 mM MgCl2, 0.05 mM EDTA, 50 ng/μl Salmon Sperm DNA (Sigma Aldrich) and 9.00% Ficoll 400 (Sigma Aldrich). The competition assay was performed by adding 9 pmol of unlabeled probes (450-fold excess) to the binding reaction. The reaction mixtures were incubated at 22 °C for 30 min and run on a non-denaturing 6% polyacrylamide gel in 0.5X TBE (Eurofins).

Labeling reactions were carried out with LightShift^TM^ Chemiluminiscent EMSA Kit (ThermoFisher Scientific) according to the provider instructions with the following modifications: after electrophoresis, gels were blotted overnight in 20X SSC using the TurboBlotter Kit (GE Healthcare Life Sciences) and cross-linking of the probe to the membrane UV-light was performed at 254 nm and 120 mJ/cm2 (UV stratalinker^®^ 2400, STRATAGENE). Chemiluminescence stained membranes were exposed to a CDD camera (FUJIFILM, LAS-4000) for 50x 10 sec exposition time increments. The last images were used for quantification and were never saturated according to LAS 4000 software. To quantify the binding affinity of Abd-B-HD to the probes, the fractional occupancy (ratio of bound/ (free+bound) probe) was calculated for three replicate experiments (Fig. S12. E) using the intensity values of the bands measured in Image [35]. The mean fractional occupancy was significantly lower with *D. santomea* probes than with *D. yakuba* probes (ANCOVA, F(1,15)=10.58, p = 0.005). We found that ABD-B-HD binds both *D. yakuba* and *D. santomea* DNA (Fig. S12. B-C). ABD-B-HD binding to the *D. yakub*a probe always resulted in a stronger shift than to the *D. santomea* probe. Furthermore, the *D. santomea* cold probe did not compete as efficiently as the *D. yakuba* cold probe to prevent formation of the *D. yakuba* DNA-ABD-B-HD complex (U-test, p=0.05).

#### Abd-B RNAi and clonal analysis

To test whether *Abd-B* is required for hypandrial bristle development, we reduced *Abd-B* expression using either genetic mutations or RNAi. Two *UAS.Abd-B-RNAi lines (#51167* and *#26746*) were crossed with 3 different *GAL4* lines, *GMR18C05-GAL4*, *NP5130-GAL4* and *NP6333-GAL4*. Crosses were kept at 29 °C and the hypandrium phenotype was examined in 10-50 F1 males (Table S13). Using the genitalia *GAL4* drivers *esg-GAL4^NP5130^* [51] and *NP6333 [52]* to express *Abd-B.RNAi^51167^*, we obtained 20 males out of 100 with developed hypandrium, among which two aberrant hypandrial bristle phenotypes were found, either bristle size reduction or bristle loss (Fig. S11. A-F, n=9/11 for *NP5130*, n=8/9 for *NP6333*, Table S13). Smaller bristles might arise from a delay in *sc* expression during development [53]. Since *Abd-B* null mutations are lethal [54], we produced mitotic mutant clones for two null mutations, *Abd-B^M1^* [54] and *Abd-B^D18^* [55]. *Abd-B* mutant mitotic recombinant clones were induced by the FLP/FRT system [56] using *Abd-B^M1^* and *Abd-B^D18^* null mutations. To induce clones, ten *yw hsflp122; FRT82B hs-CD2 y^+^ M(3) w^123^*/TM2 virgin females were crossed to ten *y; FRT82B Abd-B^M1^ red[1] e[11] ro[1] ca[1]/TM6B* or *y; FRT82B Abd-B^D18^/TM3* males (stocks were kindly provided by Ernesto Sánchez-Herrero). Crosses were flipped every 24 hours and F1 progeny were heat-shocked at 38 °C for 1 hour at different stages of larval development: 24-48, 48-72, 72-98 and 96-120 hours after egg laying [57]. From both crosses, a total of 82 F1 males (Table S13) with the genotype of *yw hsflp122; FRT82B hs-CD2 y^+^ M(3) w^123^*/*FRT82B Abd-B^M1^ red[1] e[11] ro[1] ca[1]* and *yw hsflp122; FRT82B hs-CD2 y^+^ M(3) w^123^*/*FRT82B Abd-B^D18^* were examined. Hypandria were mounted and bristle clones were screened as described above. Most of the resulting males showed extreme transformation of the genitalia (Fig. S11. G-H, O-P) but 12 males out of 82 had analyzable hypandrium (twelve males for *Abd-B^M1^* and two males for *Abd-B^D18^*). Among them, 6 males were devoid of one or both hypandrial bristles (Table S13, Fig. S11. I-N). When hypandrial bristles were present, most of them were heterozygous for the *Abd-B* mutation according to the visible markers associated with somatic recombination. Together, our results suggest that *Abd-B* is required for hypandrial bristle development.

#### Immunostaining

For leg disc stainings the larvae were fed on freshly prepared Formula 4-24^^®^^ Instant Drosophila Medium, Blue (Carolina) and staged by the presence of blue staining in their gut [58]. Larvae were chosen with the most clear gut, indicating a developmental stage of 1-6 hours before pupa formation [59]. Head parts of the larvae were cut and fixed in 4% PFA in PBS pH 7.4 for 20 minutes at room temperature. For pupal leg preparations the anterior part of the pupae were cut and fixed in 4% PFA in PBS pH 7.4 for 50 minutes at room temperature. Following fixation, samples were washed three times for 5 minutes in PBS containing 0.1% Tween20 and then permeabilized in TNT buffer (TRIS-NaCl buffer containing 0.5% Triton X-100) for 10 minutes. Samples were washed in 5% BSA in TNT for up to 5 hours at room temperature and then incubated with rabbit anti-GFP primary antibodies (Thermofisher #A6455) diluted in 1:1000 in TNT overnight at 4 °C and rinsed in TNT three times for 10 minutes at room temperature. Then, samples were washed in 5% BSA in TNT for up to 5 hours at room temperature and incubated with donkey Anti-Rabbit Dylight^^®^^ 488 (Thermofisher) secondary antibodies diluted 1:200 in TNT overnight at 4 °C. After washing the preparations in TNT for 5 minutes DNA was stained in 1μg/μl DAPI solution (Sigma-Aldrich) for 30 minutes at room temperature. The preparations were finally washed in TNT three times for 5 minutes and the imaginal discs and pupal legs were dissected in PBS and mounted in Vectashield^^®^^ H-1000. Images were acquired using Spinning Disc CSU-W1. Number of GFP-positive cells were counted in the z-stack using Image J [35] in a blind fashion regarding the genotypes using randomized file names.

**Fig. S1.**
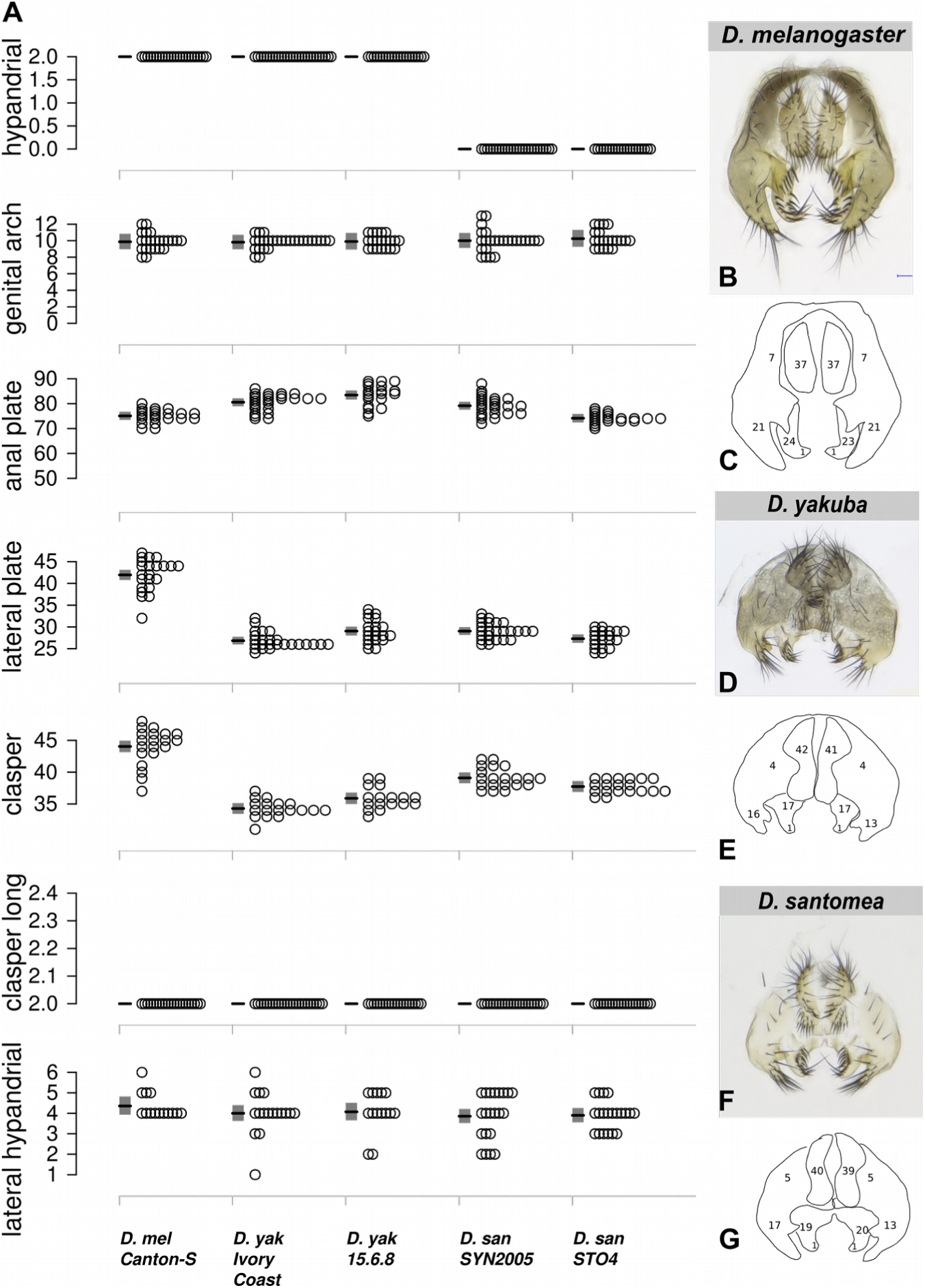
Genital bristle number in *D. yakuba* and *D. santomea*. (A) The bristle number of various bristle types is indicated for several strains of *D. melanogaster*, *D. yakuba* and *D. santomea*. Each grey dot represents one individual raised at 25°C. Mean (black line) and 95% confidence interval (grey rectangle) from a fitted GLM Quasi-Poisson model are shown. *D. yakuba* (yak) has fewer clasper bristles than *D. santomea* (san) (mean_yak= 35, mean_san = 38, GLMM-Poisson, Chisq(1) = 5. 01, p = 0.025) and more anal plate bristles than *D. santomea* (mean_yak = 82, mean_san = 77, GLMM-Poisson, Chisq(1) =3.87 p = 0.049). Dissected external genitalia from *D. melanogaster* Canton-S (B), *D. yakuba* Ivory Coast (D) and *D. santomea* SYN2005 (F) and their schematic representations (C,E,G) indicating the number of bristles in each genitalia part.

**Fig. S2.**
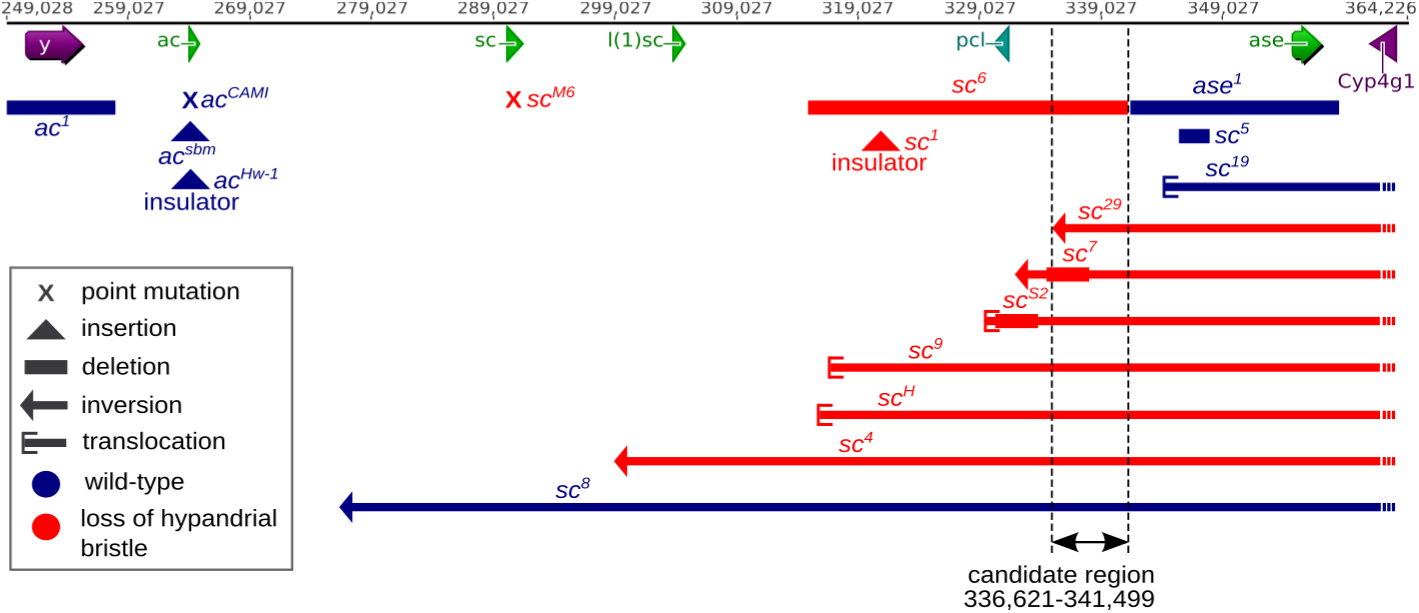
Analysis of *achaete-scute* mutants in *D. melanogaster* reveals a 5-kb candidate region driving expression in hypandrial bristles. Genomic organization of the *AS-C* locus in *D. melanogaster*. Arrows and arrowheads indicate the coding regions of *yellow* (*y*), *achaete* (*a*), *scute* (*sc*), *lethal of scute* (*l(1)sc*), *pepsinogen-like* (*pcl*), *asense* (*ase*) and *cytochrome P450-4g1* (*Cyp4g1*) genes, respectively. The *achaete-scute* mutants are represented in blue if they display a wild-type phenotype of 2 bristles and in red if they show a mutant phenotype with a reduced number of hypandrial bristles. Point mutations are represented as crosses, insertions as triangles, deletions as solid horizontal bars, inversions as thin horizontal bars terminated by an arrow, and translocations as thin horizontal bars terminated by a bracket. Comparison of all the tested *achaete-scute* mutants suggests that a 5-kb region (black arrows) located 45 kb away from *sc* 5’ coding region drives expression in the developing hypandrial bristles.

**Fig. S3.**
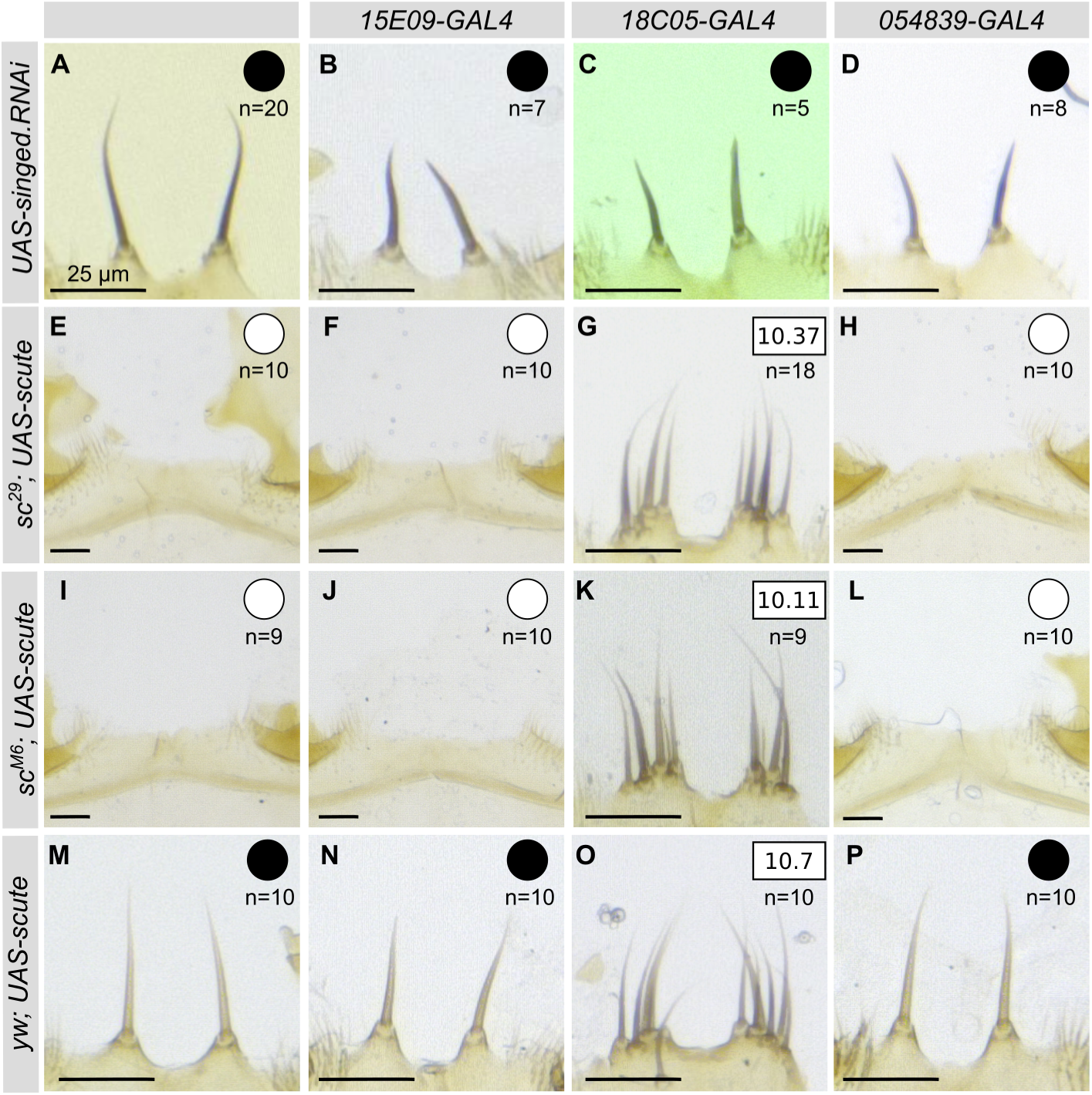
Three cis-regulatory elements of the *achaete-scute* complex (*AS-C*) govern expression in the hypandrium in *D. melanogaster*. Expression of *UAS-singed.RNAi* results in singed hypandrium bristles compared to wild-type (A) with *15E09-*(B), *18C05-* (C) and *054839-GAL4* (D). Expression of *UAS-sc* (E-P) in wild-type (M-P), *sc^29^* (E-H) or *sc^M6^* (I-L) mutant background produces extra bristles with 1*8C05-GAL4* (G,K,O) but not in absence of a *GAL4* reporter line (E,I,M) nor with *15E09-* (F,J,N) and *05439-GAL4* (H,L,P). White and black circles indicate zero or two hypandrial bristles, respectively. Average bristle numbers are shown in squares (G,K,O). n: number of scored individuals. (A-P) Scale bars indicate 25 μm.

**Fig. S4.**
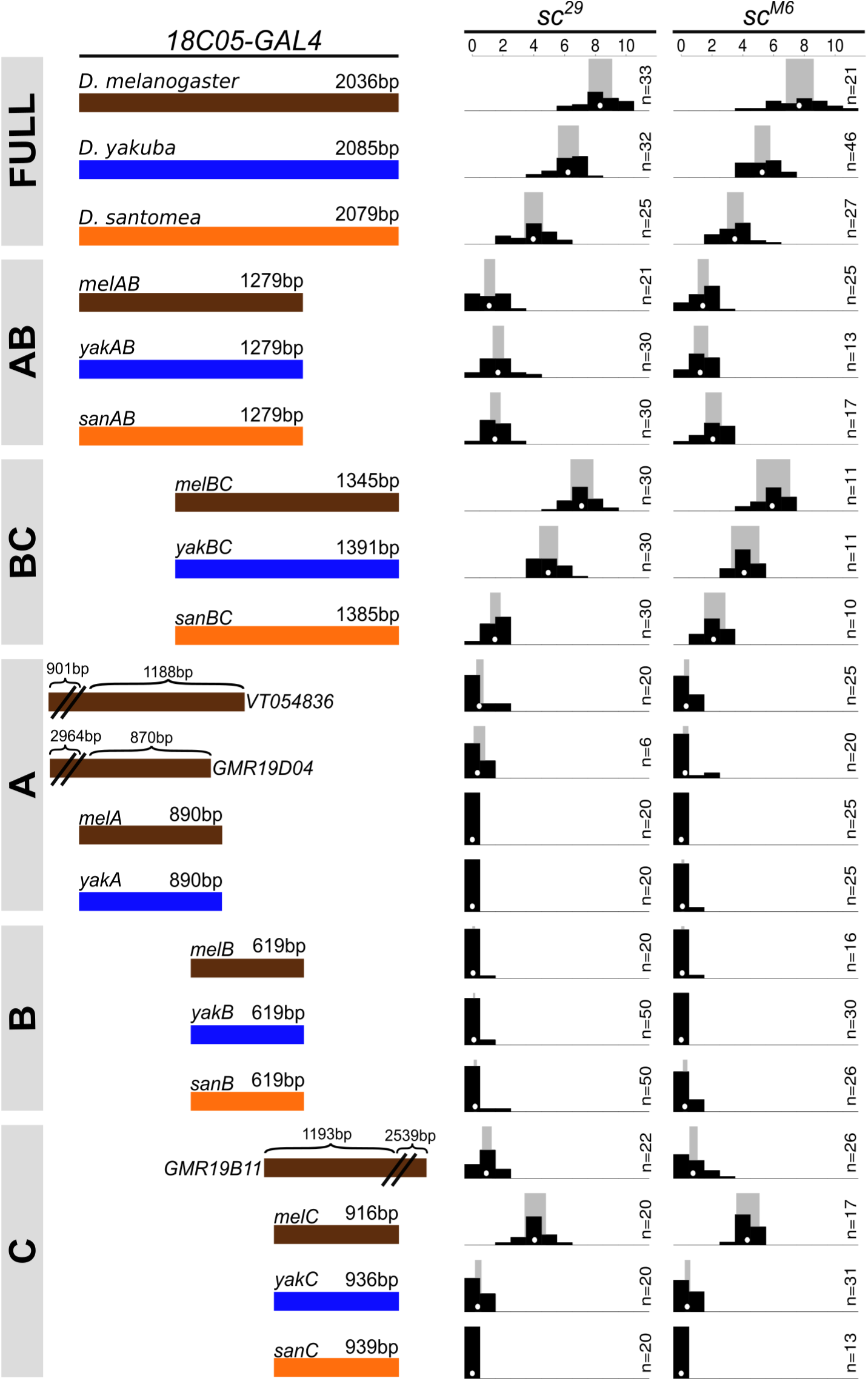
Full-length *18C05* is required for complete expression in the hypandrial bristles. Rescue of the hypandrial bristle loss of *sc^29^* (left column) and *sc^M6^* (right column) *D. melanogaster* mutants by expression of *GAL4* with *UAS-sc* driven by various genomic regions from *D. melanogaster* (brown), *D. yakuba* (blue) and *D. santomea* (orange). Distribution of hypandrial bristle number (black histogram), together with mean (white dot) and 95% confidence interval (grey rectangle) from a fitted GLM Quasi-Poisson model are shown for each genotype. n: number of scored individuals. For all three species we found that smaller segments induced significantly fewer bristles than the corresponding full region (GLM-Quasi-Poisson, F(19, 509) = 161.7, all p < 0.02 for *sc^29^*; F(19, 415) = 125.9, p < 0.03 for *sc^M6^*). Furthermore, the ~1400 bp 3’ region (region BC) from *D. santomea* rescues fewer bristles than the corresponding region from *D. melanogaster* and *D. yakuba*. Moreover, the ~900 bp 3’ region (region C) from the three species show striking differences: the region from *D. melanogaster* rescues about 4 bristles on average, the region from *D. yakuba* rescues about 0.4 bristles on average, and the region from *D. santomea* rescued no bristles. Addition of DNA 5’ of the *18C05* region increases hypandrial bristle numbers (*VT054836* different from 0 (*melA*), Exact-Poisson, p < 10-16), whereas addition of DNA 3’ of *18C05* reduces the number of bristles (*GM19B11* vs. *melC*; GLM-Quasi-Poisson, F(19, 509) = 161.7, p < 10-16 for *sc^29^*; F(19, 415) = 125.9, p < 10-16 for *sc^M6^*).

**Fig. S5.**
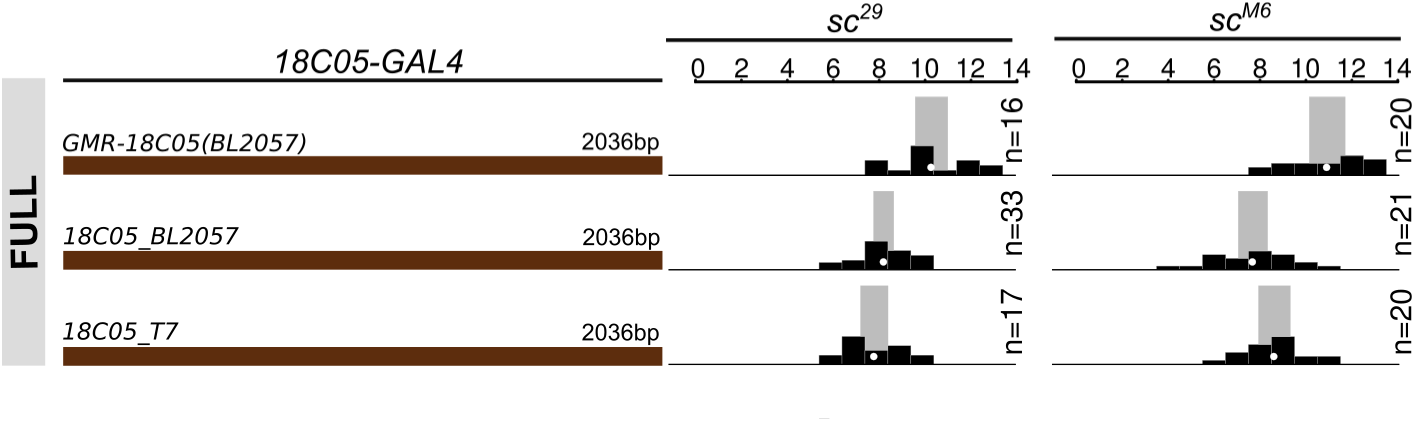
Comparison of *18C05 melanogaster-GAL4* constructs. Three different *D. melanogaster 18C05* sequences were tested with *UAS-sc* in the hypandrium in *sc^29^* (first column) and *sc^M6^* (second column). Distribution of bristle number (black histogram), together with mean (white dot) and 95% confidence interval (grey rectangle) from a fitted GLM Quasi-Poisson model are shown. Hypandrial bristle number, for *GMR-18C05* (*BL2057*) is significantly higher than *18C05_BL2057* and *18C05_T7* in both backgrounds (GLM-Quasi-Poisson, F(2, 63) = 16.88, both p < 10^-6 for *sc^29^*; F(2, 58) = 20.9, p < 10^-10 and p < 10^-5 for *sc^M6^*). *GMR-18C05* (*BL2057*) comes from the Janelia Farm collection (*35*). *18C05_BL2057* and *18C05_T7* were cloned in this study. *GMR*-*18C05* and *18C05_BL2057* are the same sequence (from *D. melanogaster* Bloomington Stock Center Strain #2057), cloned in opposite direction. *18C05_T7* contains the *18C05* sequence of *D. melanogaster T.7* strain. The *GMR-18C05* fragment is inserted in the expression vector 3’-5’ compared to the *D. melanogaster* genome sequence. In contrast, the *18C05_BL2057* and *18C05_T7* are cloned 5’-3’. Comparing bristle number between *GMR-18C05* and *18C05_BL2057* shows that the orientation of the cis-regulatory region has an effect on bristle number. All our *18C05* constructs were inserted in the same orientation, 5’-3’.

**Fig. S6.**
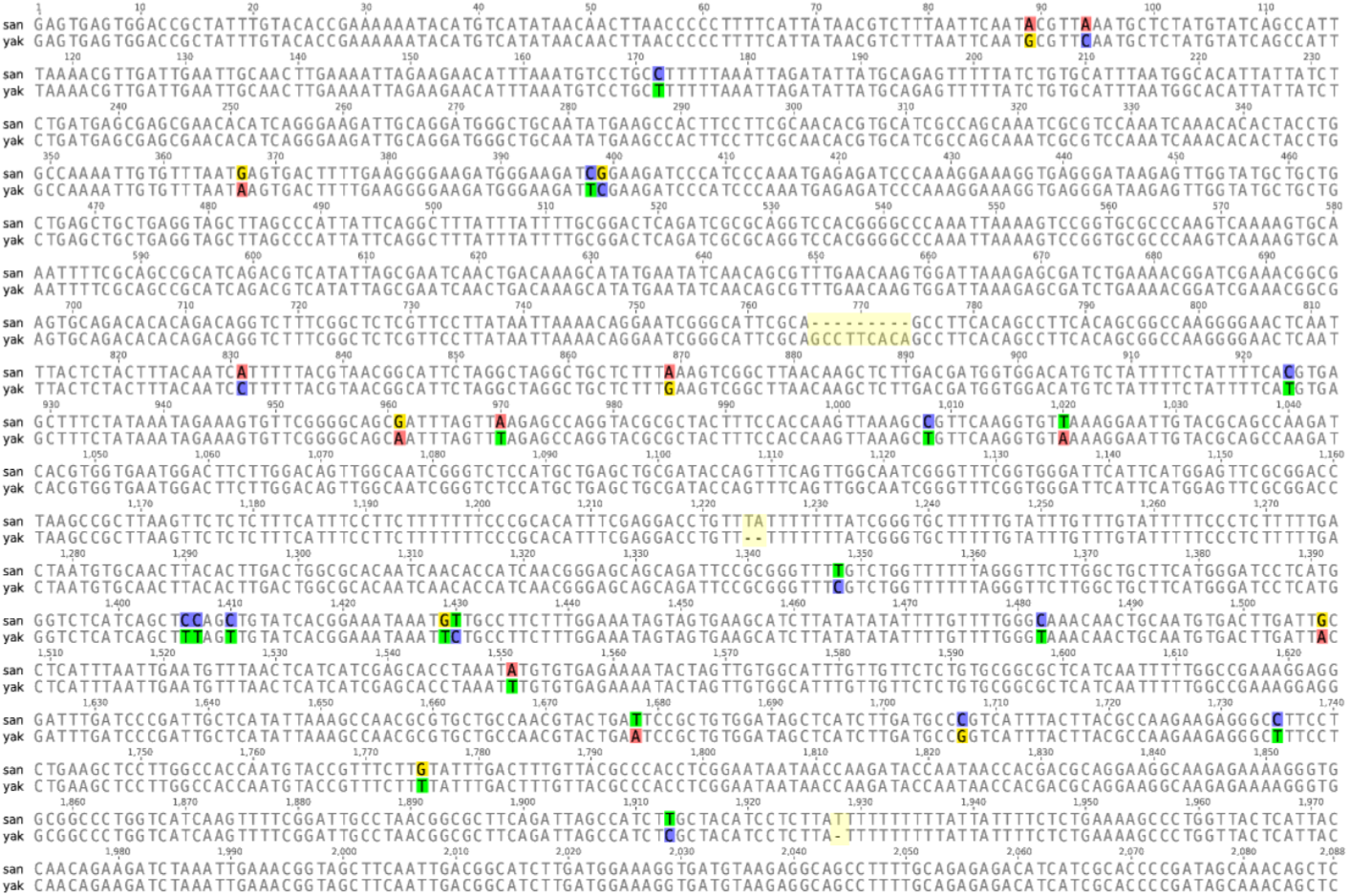
Alignment of *18C05* sequence from *D. santomea* SYN2005 (san) and *D. yakuba* Ivory Coast (yak).

**Fig. S7.**
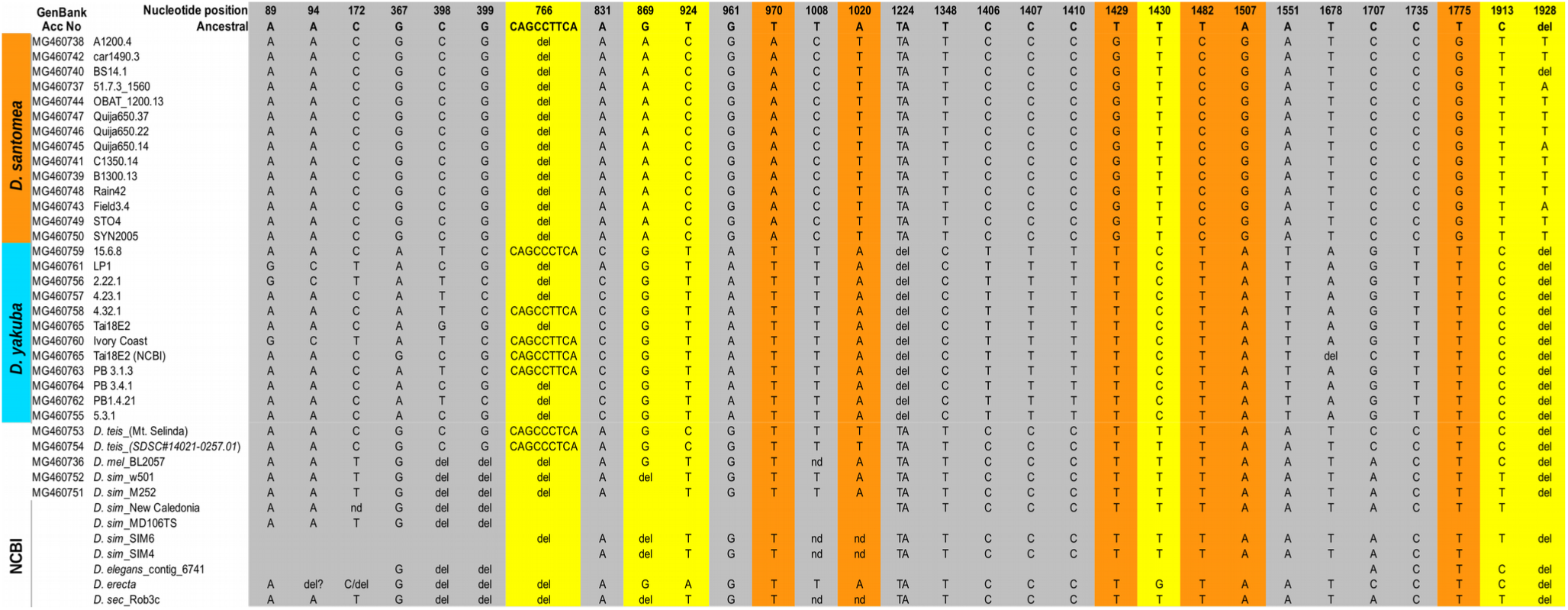
Twelve substitutions are fixed in *D. santomea 18C05*. Summary of the alignment of *18C05* sequences from different lines of 7 species. Only divergent sites within and between *D. yakuba* and *D. santomea* are shown (divergence to *D. teissieri* is not shown). Nucleotide positions are indicated in the first row. The *D. santomea*-specific substitutions are labelled in orange when they are fixed in the other *D. melanogaster* subgroup species and in yellow when they are variable in at least one of the other *D. melanogaster* subgroup species. The reconstructed ancestral sequence of *D. santomea* and *D. yakuba* is shown at the top. Del: deletion. NCBI: sequences were retrieved from NCBI BLAST. GenBank accession numbers are shown in the first column for the sequences obtained in this study.

**Fig. S8.**
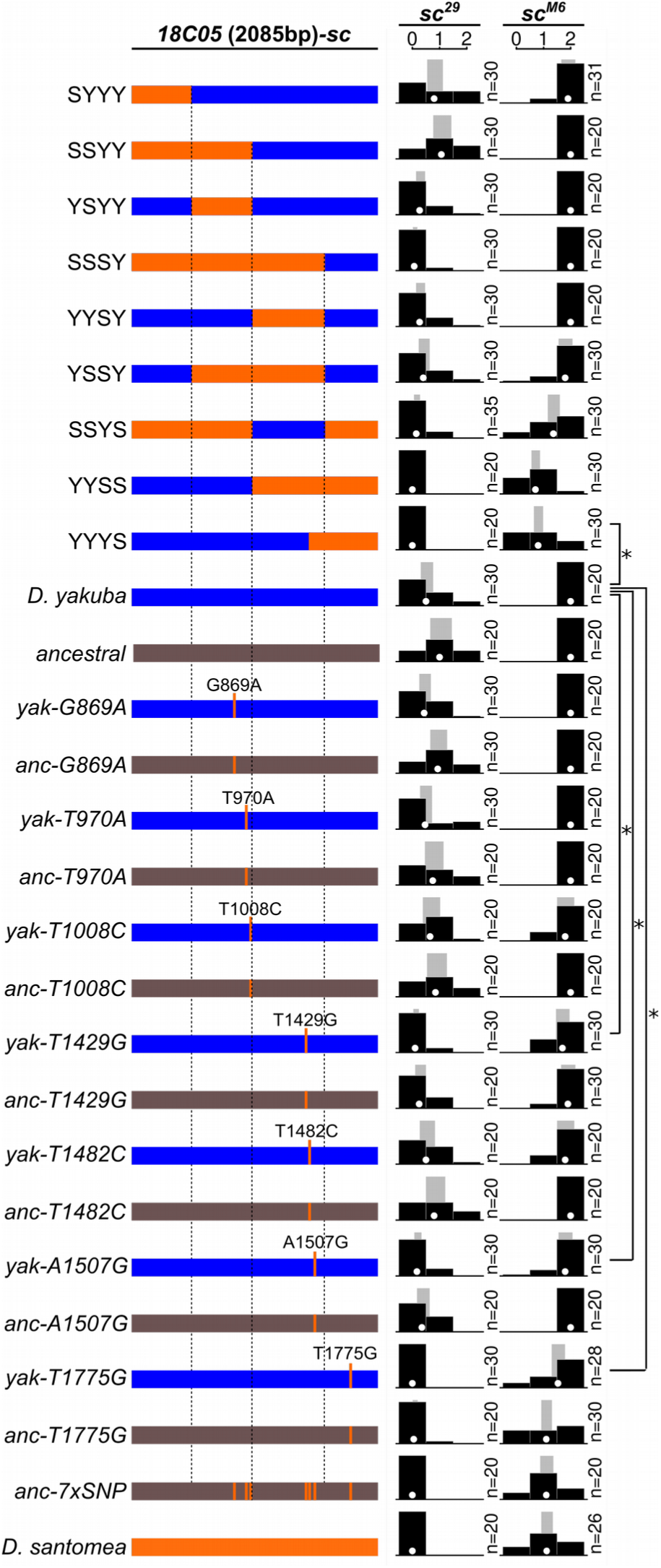
Three *D. santomea*-specific substitutions in *18C05* cause the loss of hypandrial bristles. Various full-length *18C05* regions were cloned in front of the *D. melanogaster sc* coding region and tested for their ability to rescue hypandrial bristles in *sc^29^* (left column) and *sc^M6^* (right column) mutant backgrounds. The *18C05* sequence from *D. yakuba* (blue, labeled with Y) and from *D. santomea* (orange, labeled with S) were divided into four subregions which were then fused together in different chimeric combinations. *D. santomea*-specific substitutions are represented as dark orange bars. Seven *D. santomea*-specific substitutions were introduced into either the *D. yakuba* region (blue) or the ancestrally reconstructed *18C05* region (dark grey). n: number of scored individuals. *: p<0.05. Replacing the 5’ half (1-1020 bp) of *D. yakuba 18C05* by *D. santomea* corresponding region has little or no effect (GLM-Quasi-Poisson, F(26,658) = 12.09, SSYY versus *D. yakuba*: p=0.031 in *sc^29^*, YSYY not different from *D. yakuba*: p = 0.432 in *sc^29^*, SYYY not different from *D. yakuba*: p =0.432 in *sc^29^*) whereas replacing the 3’ end (1573-2088 bp) decreases bristle number (GLM-Quasi-Poisson, F(26,618) = 16.04, YYYS versus *D. yakuba*: p<10-16 in *sc^M6^*, YYSS versus *D. yakuba*: p<10-16 in *sc^M6^*). The 3’ middle part (1021-1572 bp) has no effect (YYSY not different from *D. yakuba*: p = 0.432 in *sc^29^*, YYSS versus YYYS: p = 0.579 in *sc^M6^*) except in presence of the *D. santomea* 5’ half (1-1020 bp), in which case it decreases bristle number (SSSY versus SSYY: p<10-4 in *sc^29^*). Our analysis of chimeric constructs thus indicates that at least two changes, in regions 1021-1572 and 1573-2088, contribute to the reduced ability of *D. santomea 18C05* to produce hypandrial bristles. Three substitutions significantly decreased the number of rescued bristles in the *D. yakuba* and/or the ancestral sequence (T1429G: p=0.021 and p=0.009 in *sc^29^*, respectively; A1507G: p=0.066 and p=0.03 in *sc^29^*, respectively; T1775G: p=0.013 and p=10-8 in *sc^M6^*, respectively).

**Fig. S9.**
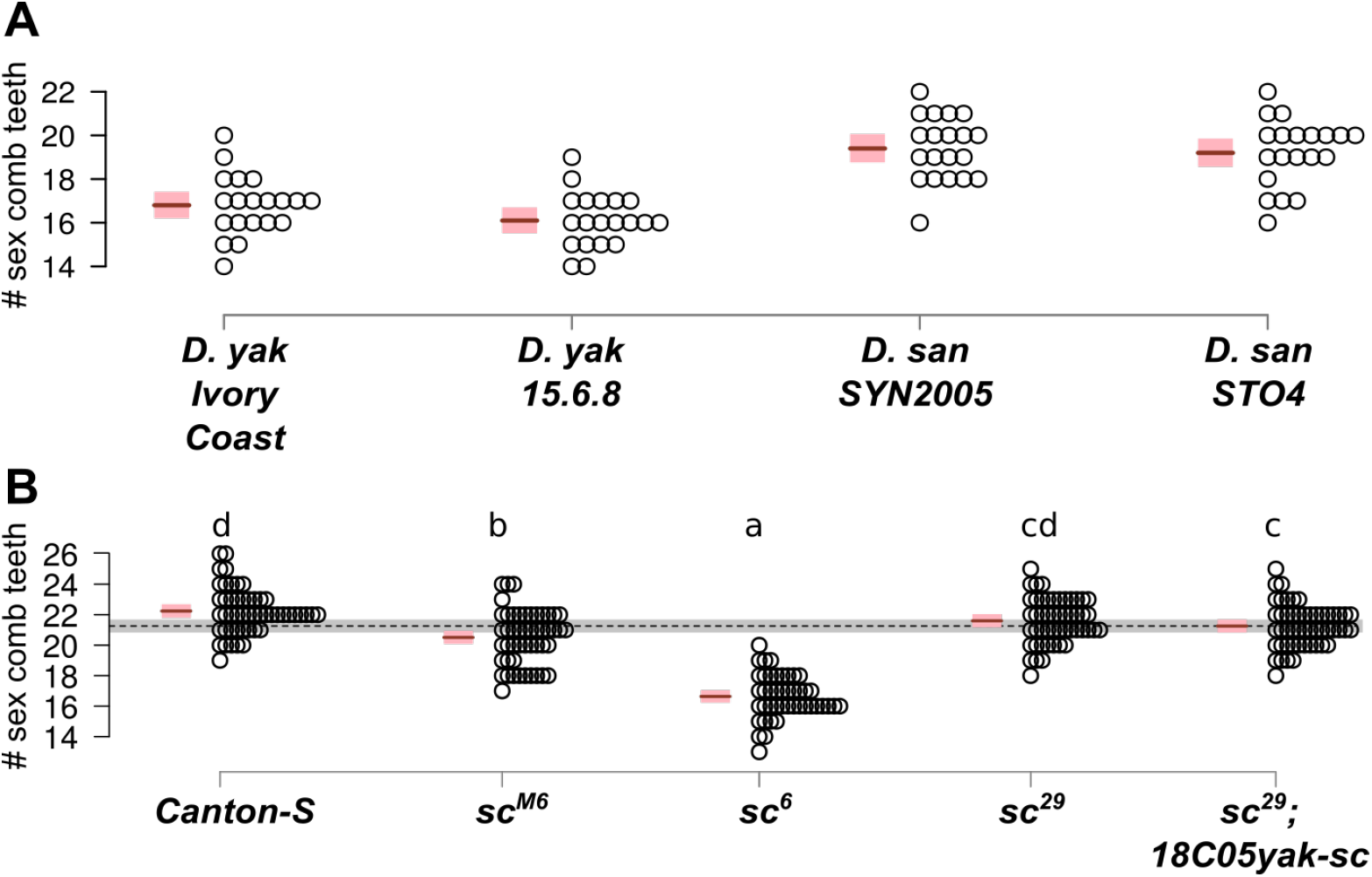
Sex comb tooth number in *D. yakuba* and *D. santomea* and *D. melanogaster scute* mutants. Each circle represents one individual raised at 25° C. Mean (brown line) and 95% confidence interval (pink rectangle) from a fitted GLM Quasi-Poisson model are shown. (A) Sex comb tooth number is significantly different between *D. yakuba* and *D. santomea* (GLM-Poisson, Chisq(1)=2.76, p = 0.007). (B) *sc^M6^* and *sc^6^* have significantly different sex comb tooth number than Canton-S. Letters indicate the results of all-pairwise comparisons after Holm-Bonferroni correction. Two genotypes are significantly different from each other (p < 0.05) when they do not share a letter. Sex comb tooth number of *sc^29^* mutant males is not significantly different from wild-type (F(4,239)=100.06, p=0.09) and the *18C05yak-sc* construct does not significantly increase sex comb tooth number in the *sc^29^* background (*sc^29^* versus *sc^29^18C05yak-sc*, F(4,239)=100.06, p=0.27), so we did not examine the effects of *D. santomea* substitutions in the *sc^29^* background.

**Fig. S10.**
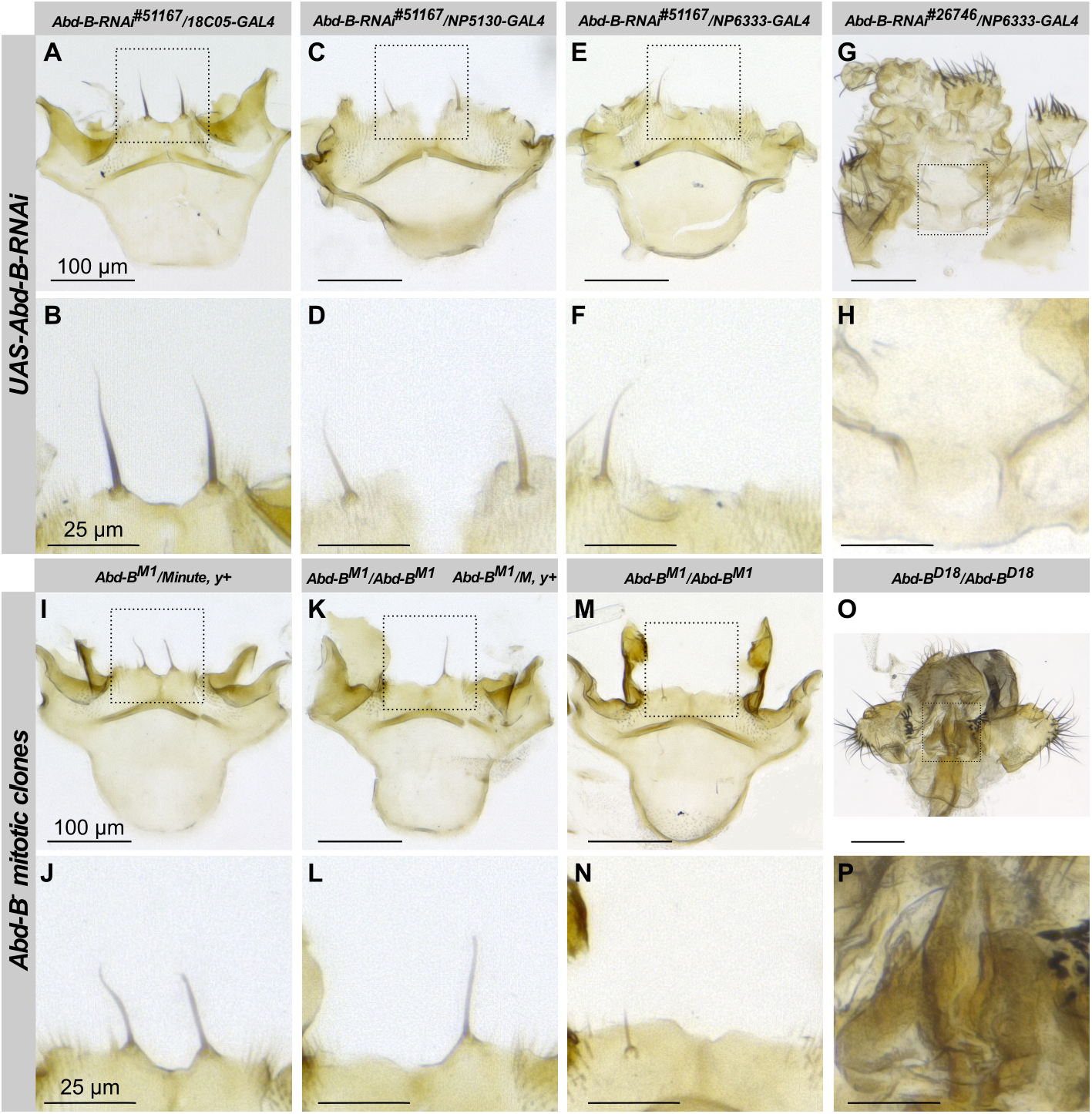
Hypandrial bristle development affected in *Abd-B-RNAi* and *Abd-B^−^ clones*. Wild-type hypandrial bristles. (B) In *Abd-B-RNAi^#51167^/GMR18C05-GAL4* males the hypandrial bristles are thiner. (C) In *Abd-B-RNAi^#51167^/NP5130-GAL4* hypandrial bristles are thiner and shorter. (D) In *Abd-B-RNAi^#51167^/NP6333-GAL4* hypandrial bristles are thiner or lost. (I-N) In *Abd-B^−^* clones hypandrial bristles are lost. Clones were selected in *Minute, y^+^* background. One hypandrium with two *Minute, y^+^* bristles (I-J), one hypandrium with only *Minute, y^+^* bristle (K-L) and one hypandrium with no hypandrial bristle (M-N) are shown. Extreme transformations of genitalia with abnormal hypandrium are shown in *Abd-B-RNAi^#26746^/NP6333-GAL4* (G-H).

**Fig. S11.**
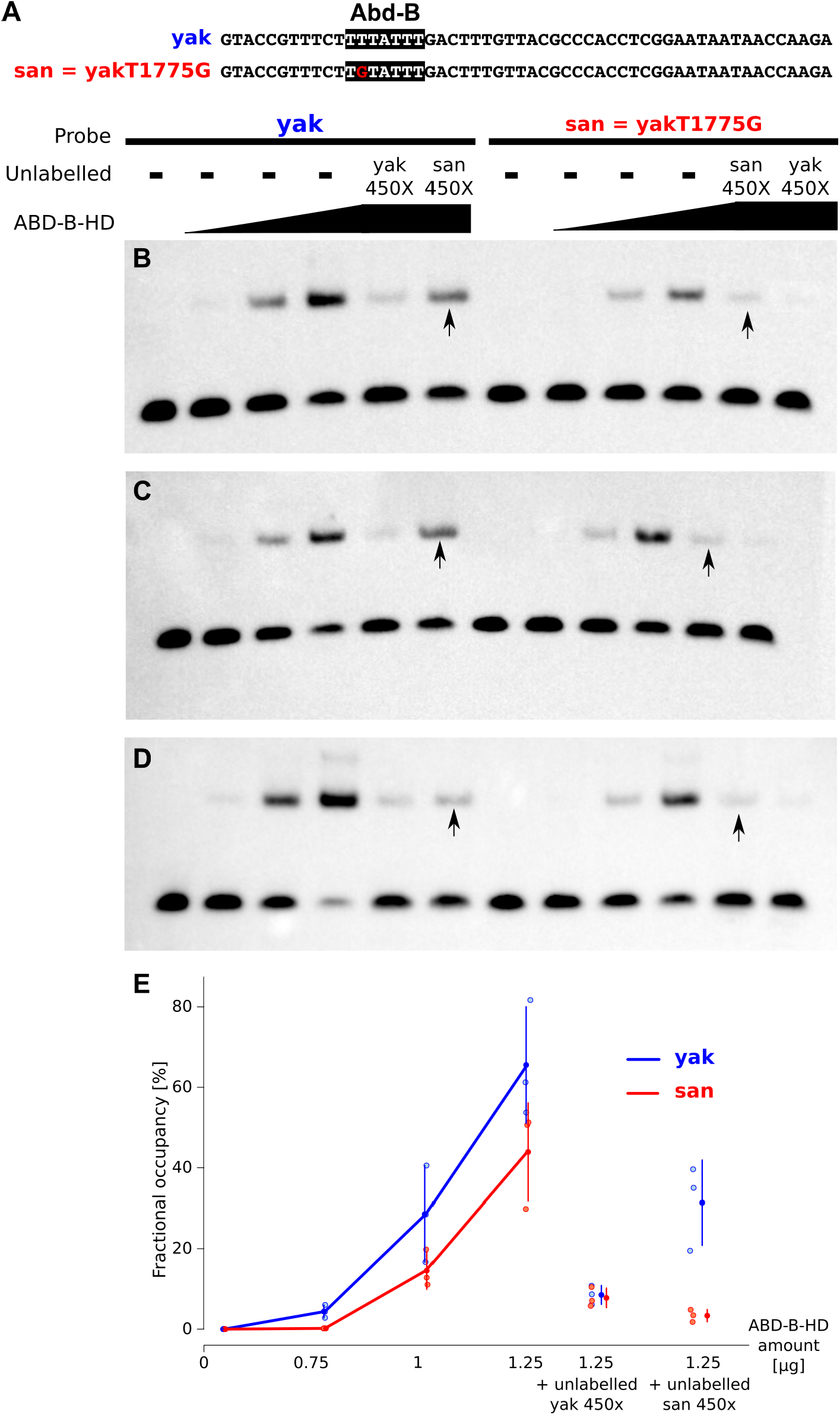
*D. santomea*-specific substitution T1775G affects Abd-B binding. (A) Two double-stranded oligonucleotides were used in EMSA, one with *D. yakuba* sequence (yak, blue) and one with *D. santomea* sequence (san=yakT1775G, red). The putative Abd-B binding site is shown in black. (B-D) Threee paralel EMSA were performed with increasing amount of Abd-B-HD (0.75 μg, 1.0 μg and 1.25 μg) and *yak* probe or *san* probe. In competitor lanes 450x excess of unlabelled *yak* probe or *san* probe were added to the binding reaction. In all experiments *san* probe binds less than yak probe to Abd-B-HD, and unlabelled *san* probe competes less the band shift of the labelled probe compared to the *yak* probe (indicated with arrows). (E) Quantification of EMSA shifts. Fractional occupancy (ratio of bound/(free+bound) probe) is shown for three parallel experiments. Mean (dot) and standard deviation (bar) are shown for *yak* and *san* probes, in blue and red, respectively.

**Table S1.**
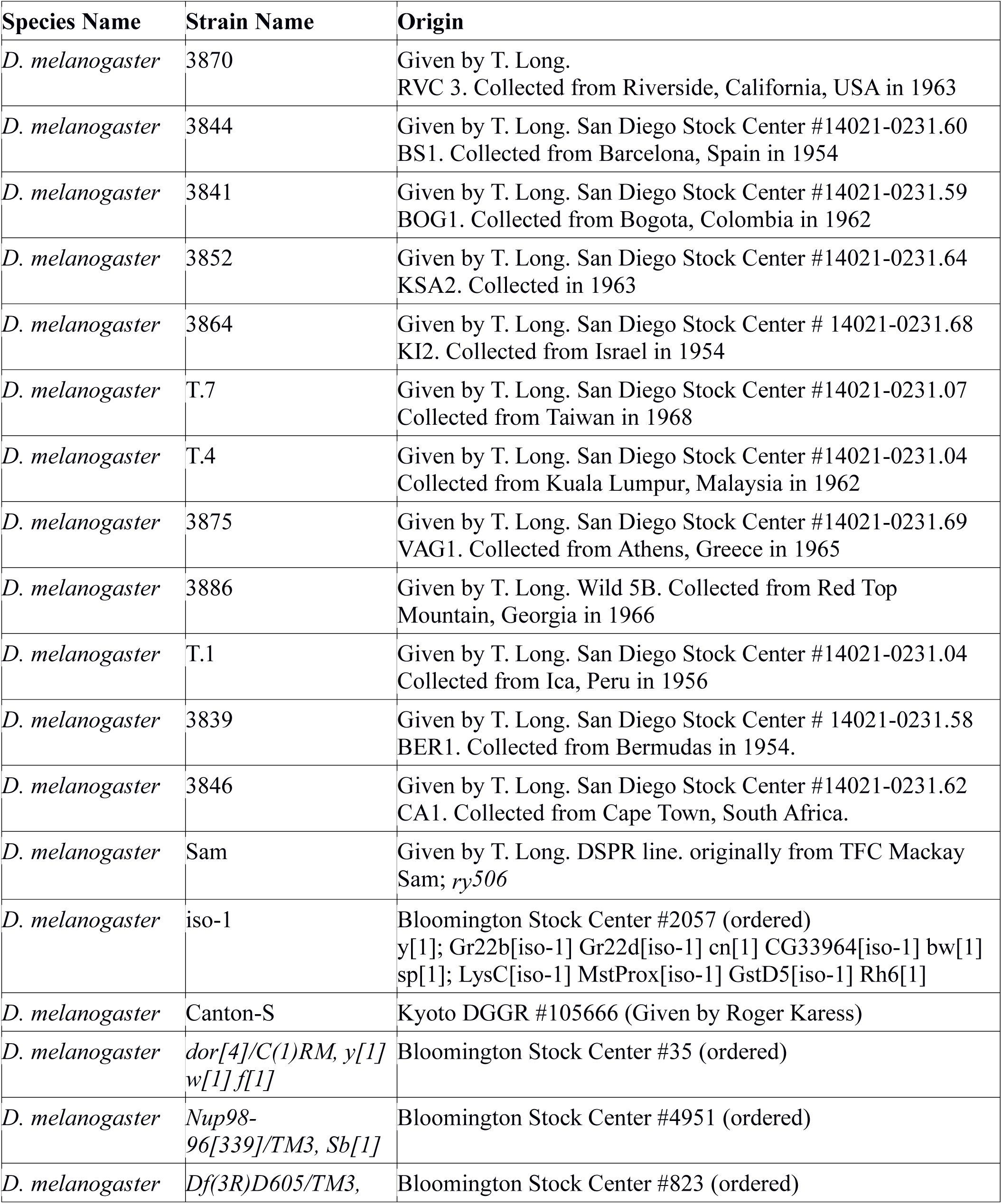
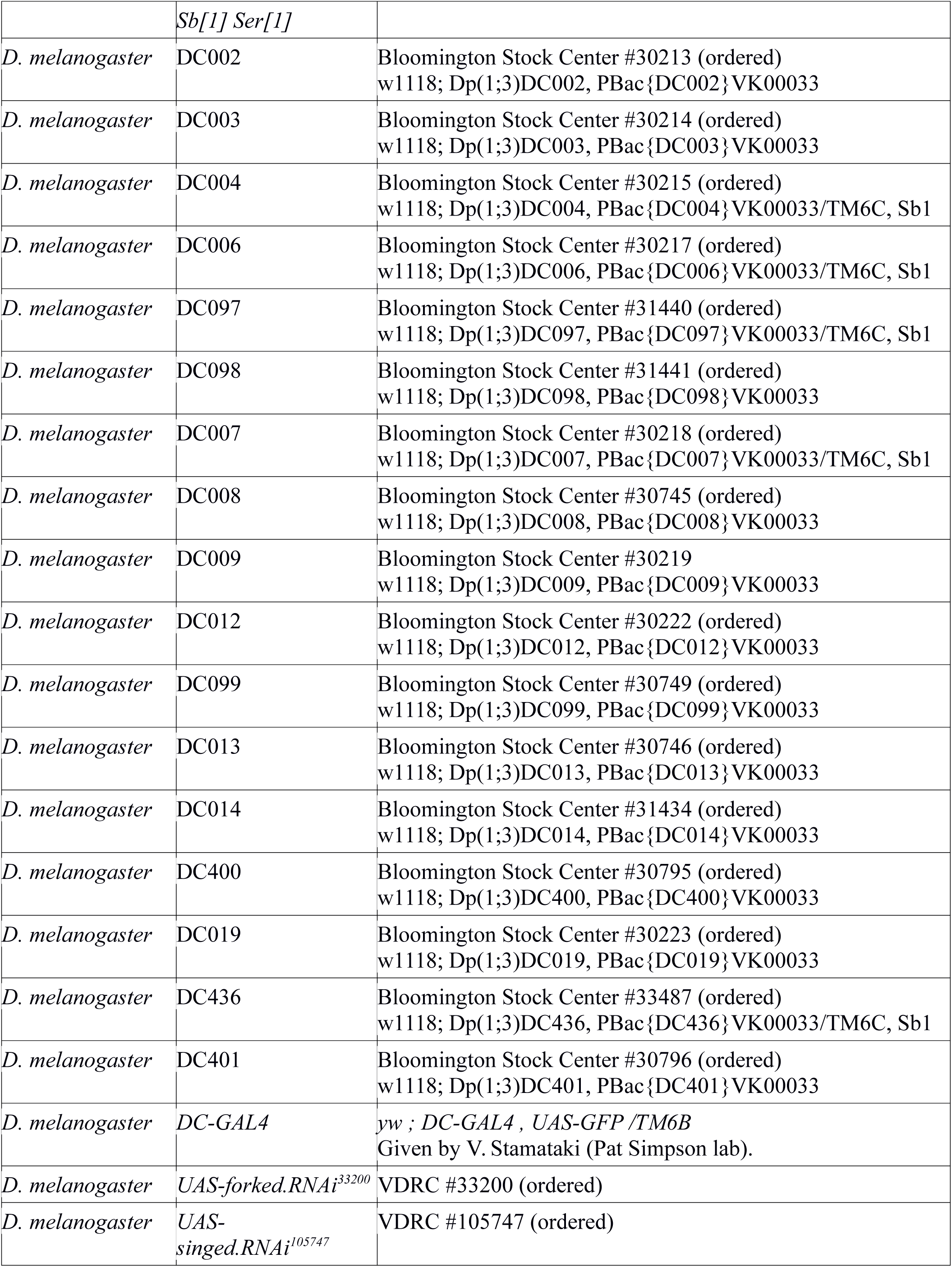
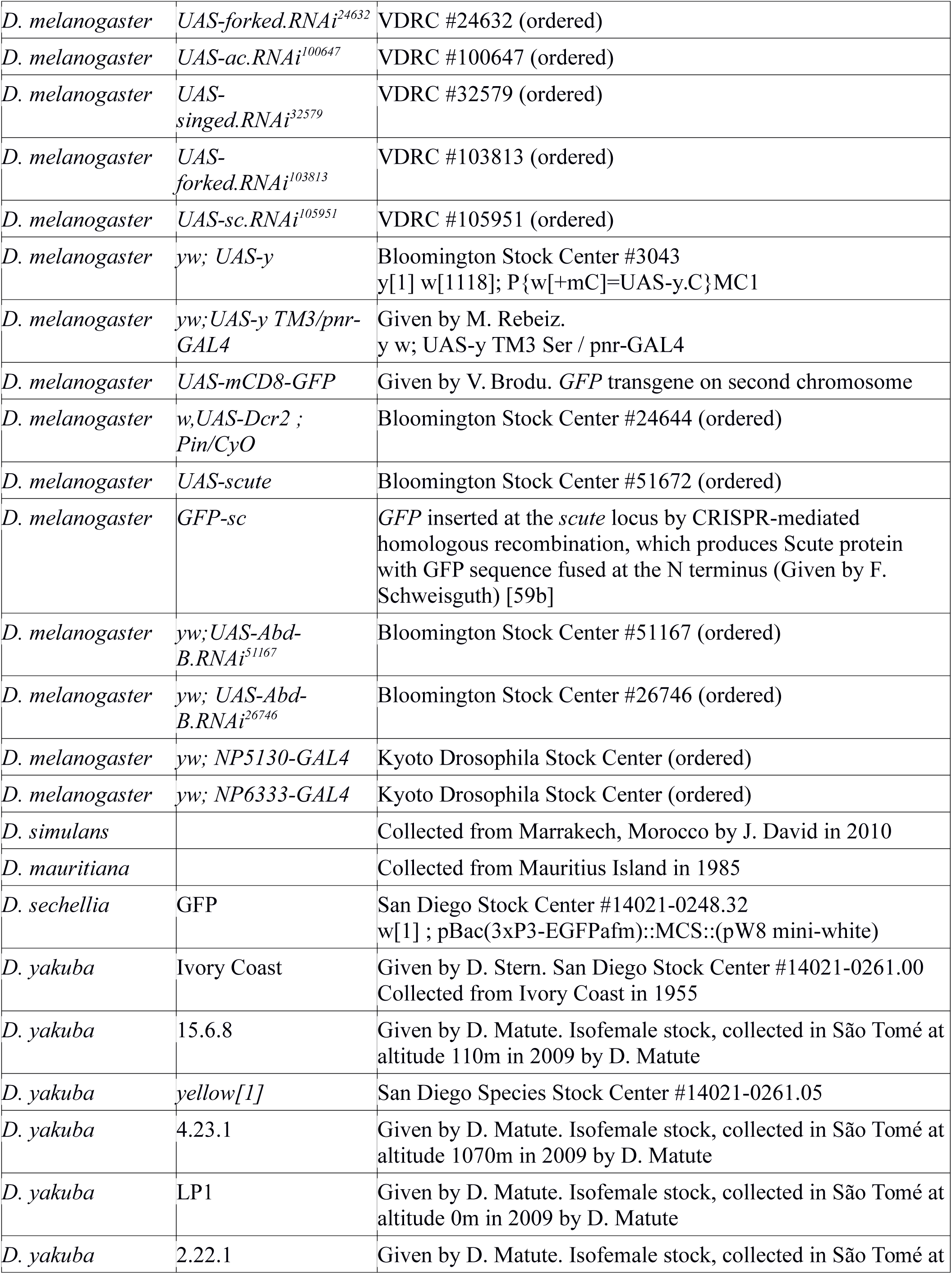
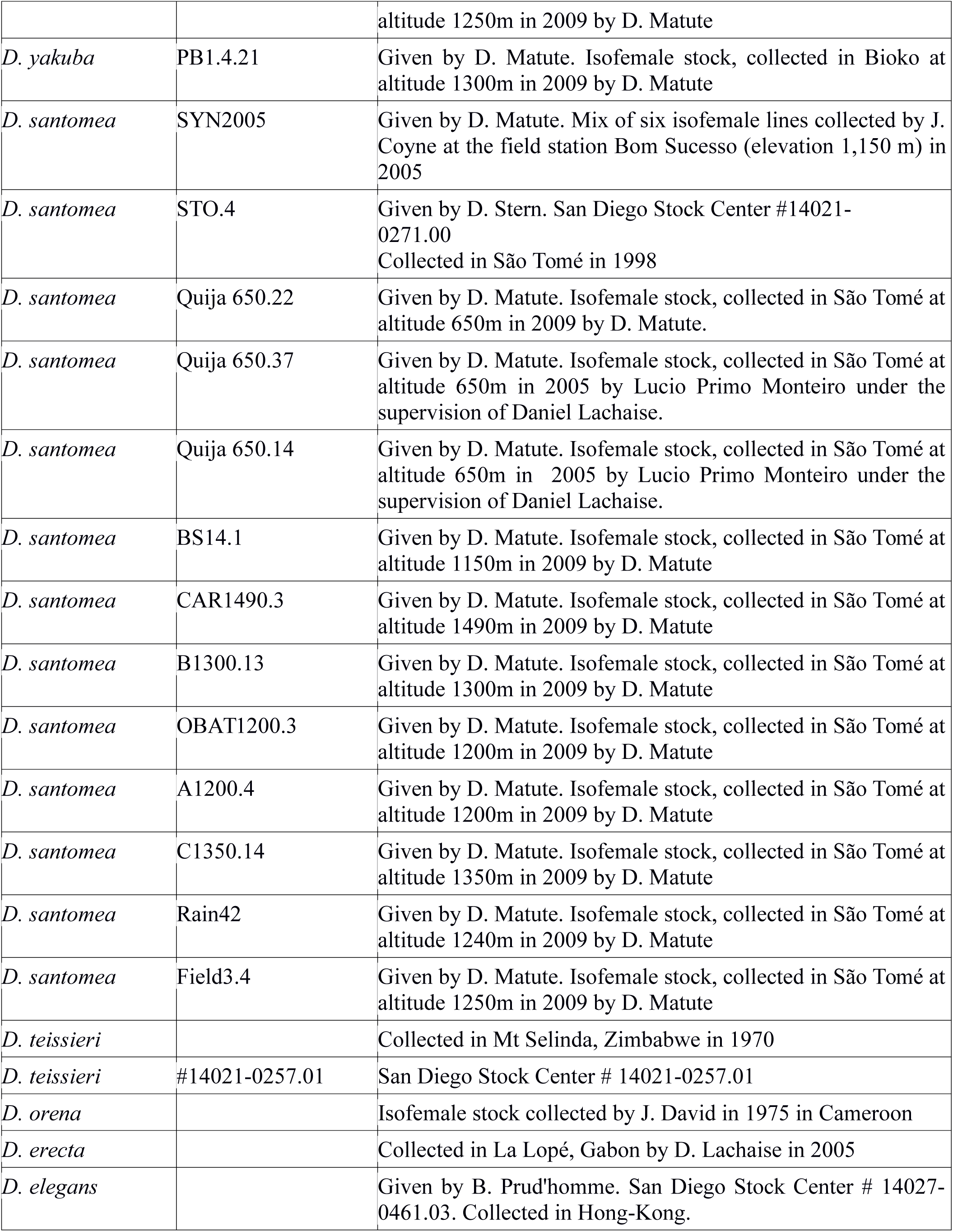
Fly lines used. See Table S4 for *GAL4* lines others than *DC-GAL4* and Table S5 for *achaete-scute* mutant lines.

**Table S2.** Hypandrial bristle number in pure species and F1 hybrids. ^0^Flies raised at 25°C,^1^Flies raised at 29°C, ^2^Flies raised at 18°C, ^3^Flies raised at 21°C, ^4^Flies raised at 14°C, ^5^Flies collected directly from the wild.

|  | Number of Flies with 0 bristles | Number of Flies with 1 bristles | Number of Flies with 2 bristles | Number of Flies with 3 bristles | Total Number of Flies |
| --- | --- | --- | --- | --- | --- |
| ***D. melanogaster*** |  |  |  |  |  |
| 3870<sup>0</sup> | 0 | 0 | 20 | 0 | 20 |
| 3844<sup>0</sup> | 0 | 0 | 21 | 0 | 21 |
| 3841<sup>0</sup> | 0 | 0 | 21 | 0 | 21 |
| 3852<sup>0</sup> | 0 | 0 | 21 | 0 | 21 |
| 3864<sup>0</sup> | 0 | 0 | 22 | 0 | 22 |
| T.7<sup>0</sup> | 0 | 0 | 20 | 0 | 20 |
| T.4<sup>0</sup> | 0 | 0 | 19 | 1 | 20 |
| 3875<sup>0</sup> | 0 | 0 | 26 | 0 | 26 |
| 3886<sup>0</sup> | 0 | 0 | 19 | 1 | 20 |
| T.1<sup>0</sup> | 0 | 0 | 19 | 1 | 20 |
| 3839<sup>0</sup> | 0 | 0 | 20 | 0 | 20 |
| 3846<sup>0</sup> | 0 | 0 | 20 | 0 | 20 |
| Sam<sup>0</sup> | 0 | 0 | 20 | 0 | 20 |
| iso-1<sup>0</sup> | 0 | 6 | 7 | 1 | 15 |
| Canton-S<sup>0</sup> | 0 | 0 | 20 | 0 | 20 |
| ***D. sechellia* GFP<sup>0</sup>** | 0 | 0 | 31 | 0 | 31 |
| ***D. mauritiana*<sup>0</sup>** | 0 | 0 | 34 | 0 | 34 |
| ***D. mauritiana*<sup>2</sup>** | 0 | 0 | 34 | 0 | 34 |
| ***D. simulans*<sup>5</sup>** | 0 | 0 | 10 | 0 | 10 |
| ***D. yakuba*** |  |  |  |  |  |
| Ivory Coast<sup>0</sup> | 0 | 0 | 32 | 0 | 32 |
| Ivory Coast<sup>2</sup> | 0 | 0 | 30 | 0 | 30 |
| 4.23.1<sup>1</sup> | 0 | 0 | 20 | 0 | 20 |
| PB1.4.21<sup>1</sup> | 0 | 0 | 25 | 0 | 25 |
| LP1<sup>1</sup> | 0 | 0 | 35 | 0 | 35 |
| 2.22.1<sup>1</sup> | 0 | 0 | 13 | 0 | 13 |
| 15.6.8<sup>1</sup> | 0 | 0 | 10 | 0 | 10 |
| ***D. santomea*** |  |  |  |  |  |
| SYN2005<sup>0</sup> | 60 | 0 | 0 | 0 | 60 |
| SYN2005<sup>2</sup> | 30 | 0 | 0 | 0 | 30 |
| STO.4<sup>0</sup> | 29 | 0 | 0 | 0 | 29 |
| Quija 650.37<sup>0</sup> | 30 | 0 | 0 | 0 | 30 |
| Quija 650.14<sup>0</sup> | 35 | 0 | 0 | 0 | 35 |
| BS14.1<sup>2</sup> | 22 | 0 | 0 | 0 | 22 |
| CAR1490.3<sup>2</sup> | 21 | 0 | 0 | 0 | 21 |
| B1300.13<sup>2</sup> | 20 | 0 | 0 | 0 | 20 |
| OBAT1200.3<sup>2</sup> | 22 | 0 | 0 | 0 | 22 |
| A1200.4<sup>2</sup> | 20 | 0 | 0 | 0 | 20 |
| C1350.14<sup>2</sup> | 20 | 0 | 0 | 0 | 20 |
| Rain42<sup>2</sup> | 23 | 0 | 0 | 0 | 23 |
| Field3.4<sup>2</sup> | 21 | 0 | 0 | 0 | 21 |
| ***D. teissieri*<sup>0</sup>** | 0 | 0 | 30 | 0 | 30 |
| *D. teissieri*<sup>2</sup> | 0 | 0 | 30 | 0 | 30 |
| ***D. orena*<sup>3</sup>** | 0 | 0 | 5 | 0 | 5 |
| ***D. erecta*<sup>4</sup>** | 0 | 0 | 10 | 0 | 10 |
| ***D. elegans*<sup>0</sup>** | 0 | 0 | 17 | 0 | 17 |
| *D. santomea*/*D. yakuba* F1 hybrids<sup>0</sup><br>(*D. santomea* females x *D. yakuba* males) | 0 | 2 | 32 | 0 | 34 |
| *D. santomea*/*D. yakuba* F1 hybrids<sup>0</sup><br>(*D. yakuba* females x *D. santomea* males) | 29 | 0 | 0 | 0 | 29 |
| ***D. melanogaster*/*D. santomea* hybrids** |  |  |  |  |  |
| san/DC002 (1A1-1A1) | 19 | 4 | 1 | 0 | 24 |
| san/DC003 (1A1-1A1) | 15 | 7 | 2 | 0 | 24 |
| san/DC004 (1A1-1A3) | 18 | 6 | 0 | 0 | 24 |
| san/DC006 (1A3-1A8) | 18 | 3 | 3 | 0 | 24 |
| san/DC097 (1A7-1B4) | 8 | 9 | 7 | 0 | 24 |
| san/DC098 (1B2-1B8) | 19 | 3 | 2 | 0 | 24 |
| san/DC007 (1B1-1B5) | 18 | 5 | 1 | 0 | 24 |
| san/DC008 (1B4-1B10) | 21 | 2 | 1 | 0 | 24 |
| san/DC009 (1B9-1B13) | 18 | 4 | 2 | 0 | 24 |
| san/DC012 (1C3-1C5) | 17 | 5 | 2 | 0 | 24 |
| san/DC099 (1C3-1C4) | 21 | 3 | 0 | 0 | 24 |
| san/DC013 (1C4-1D2) | 16 | 8 | 0 | 0 | 24 |
| san/DC014 (1D1-1D2) | 20 | 4 | 0 | 0 | 24 |
| san/DC400 (1D2-1E1) | 19 | 3 | 2 | 0 | 24 |
| san/DC019 (1E1-1E3) | 18 | 5 | 1 | 0 | 24 |
| san/DC436 (1E3-1E5) | 22 | 0 | 2 | 0 | 24 |
| san/DC401 (2A2-2B1) | 21 | 3 | 0 | 0 | 24 |
| san/TM3 (control) | 68 | 18 | 2 | 0 | 95 |

**Table S3.**
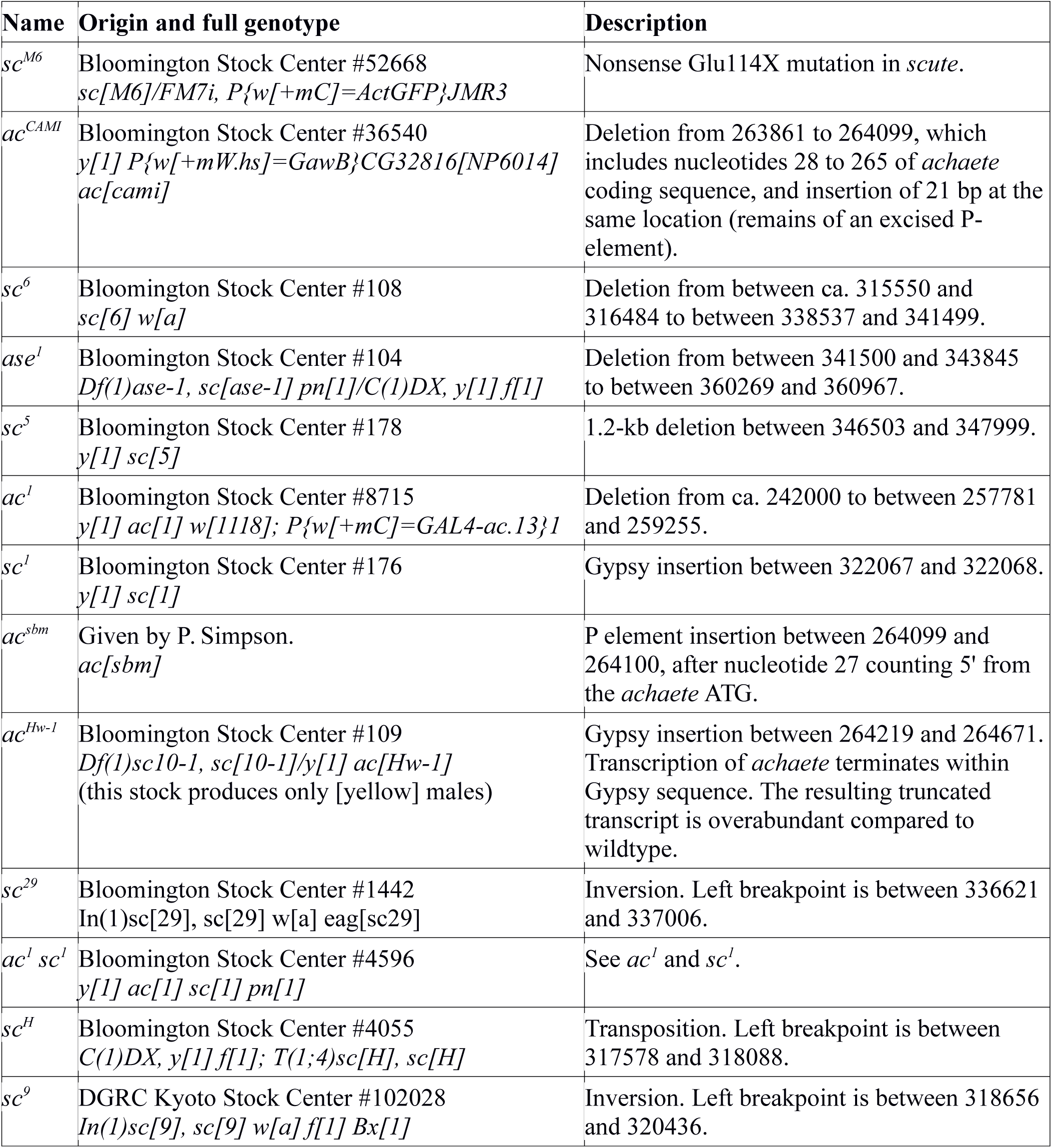
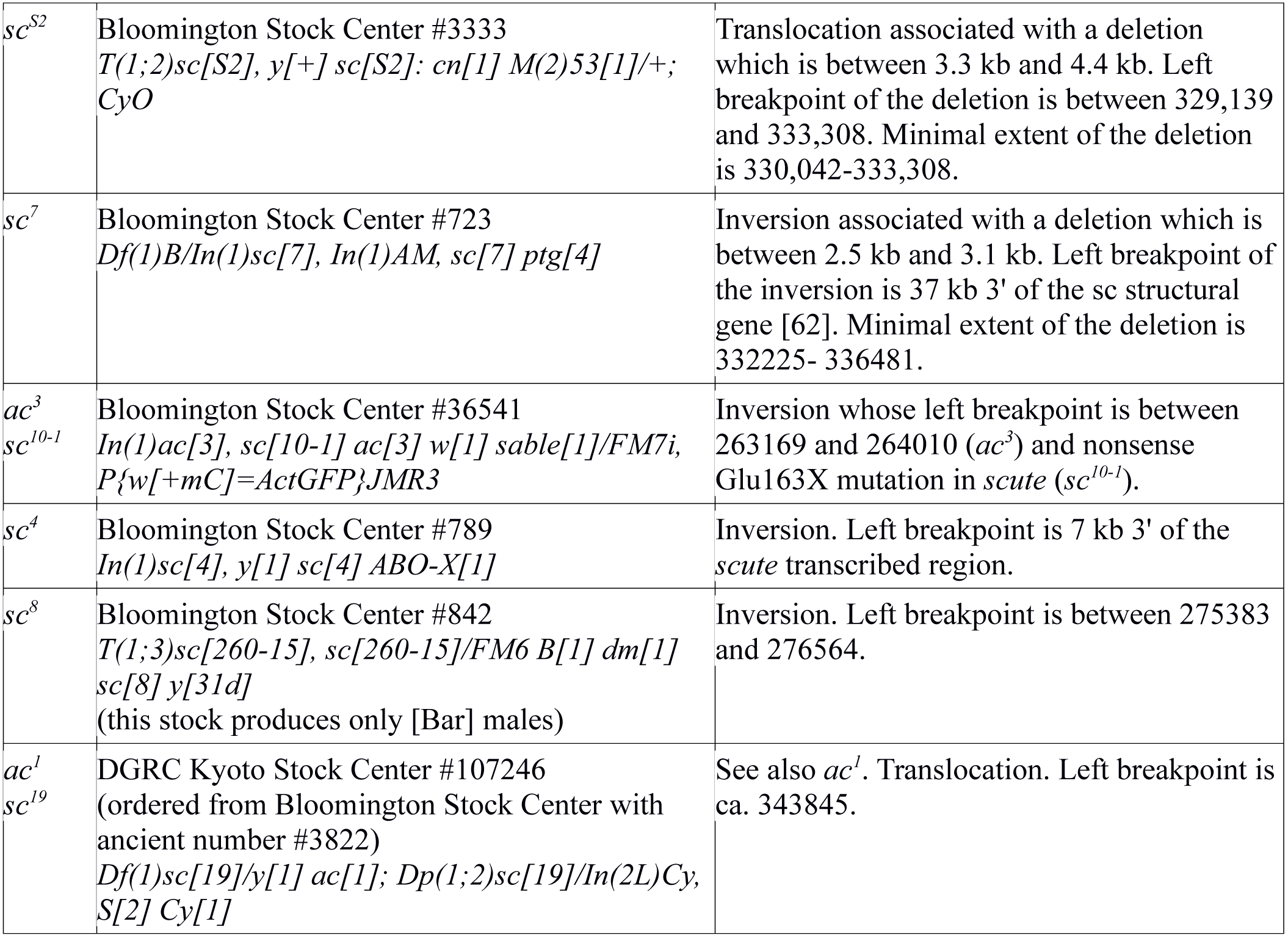
*achaete-scute* mutant lines used. Coordinates are for *D. melanogaster* reference genome iso-1 (FB2013_03, version 3). Based on restriction sites we estimated that point 0 of [60,61] corresponds to position 330342 in iso-1. Compared to other *D. melanogaster* strains, iso-1 contains a 6127-bp transposable element named 3S18{@4/TE19523 at position 322,507-328,633. Coordinates were adjusted to account for the shift due to the transposable element.

**Table S4.**
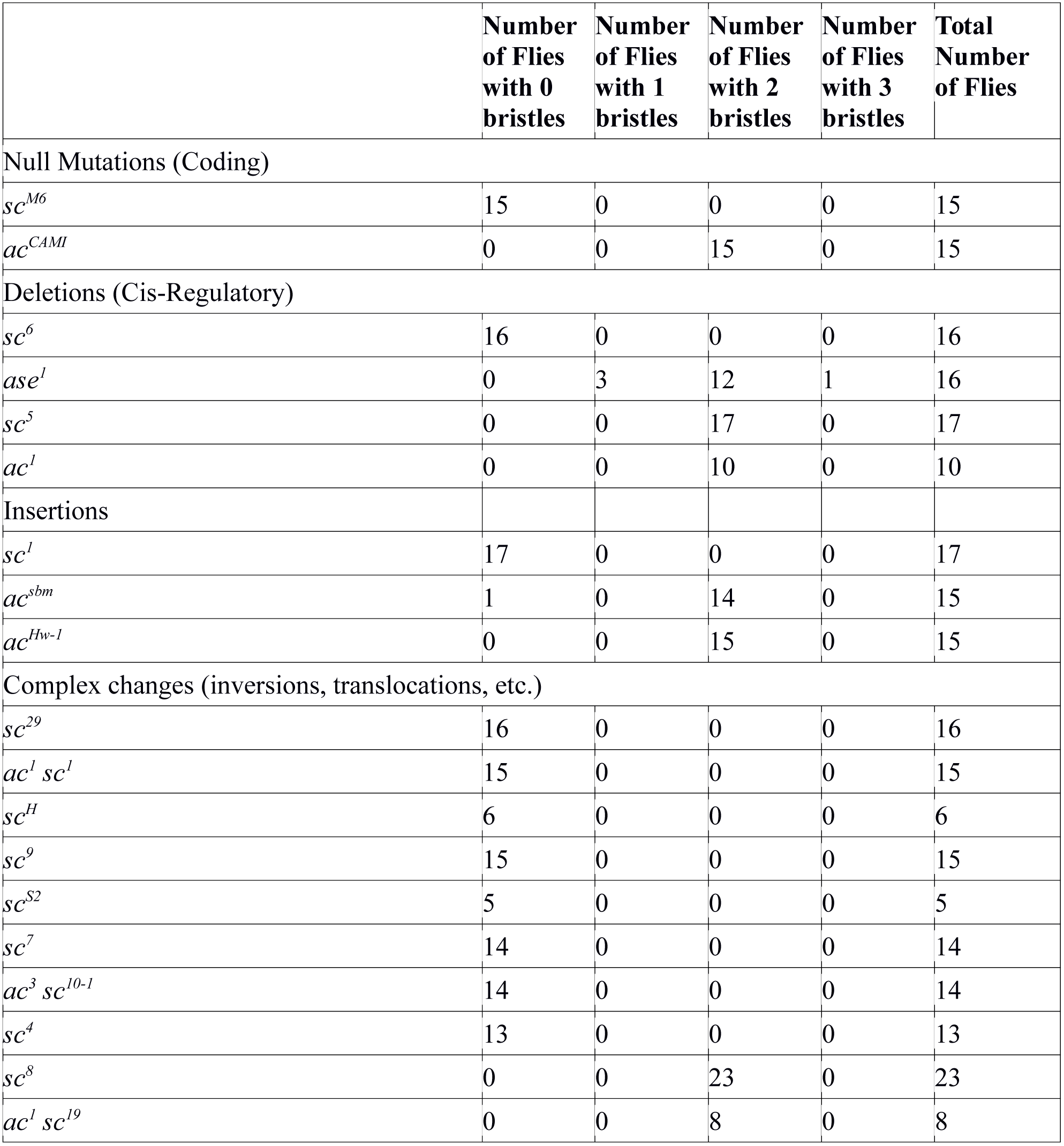
Hypandrial bristle number in *achaete-scute* mutants

**Table S5.**
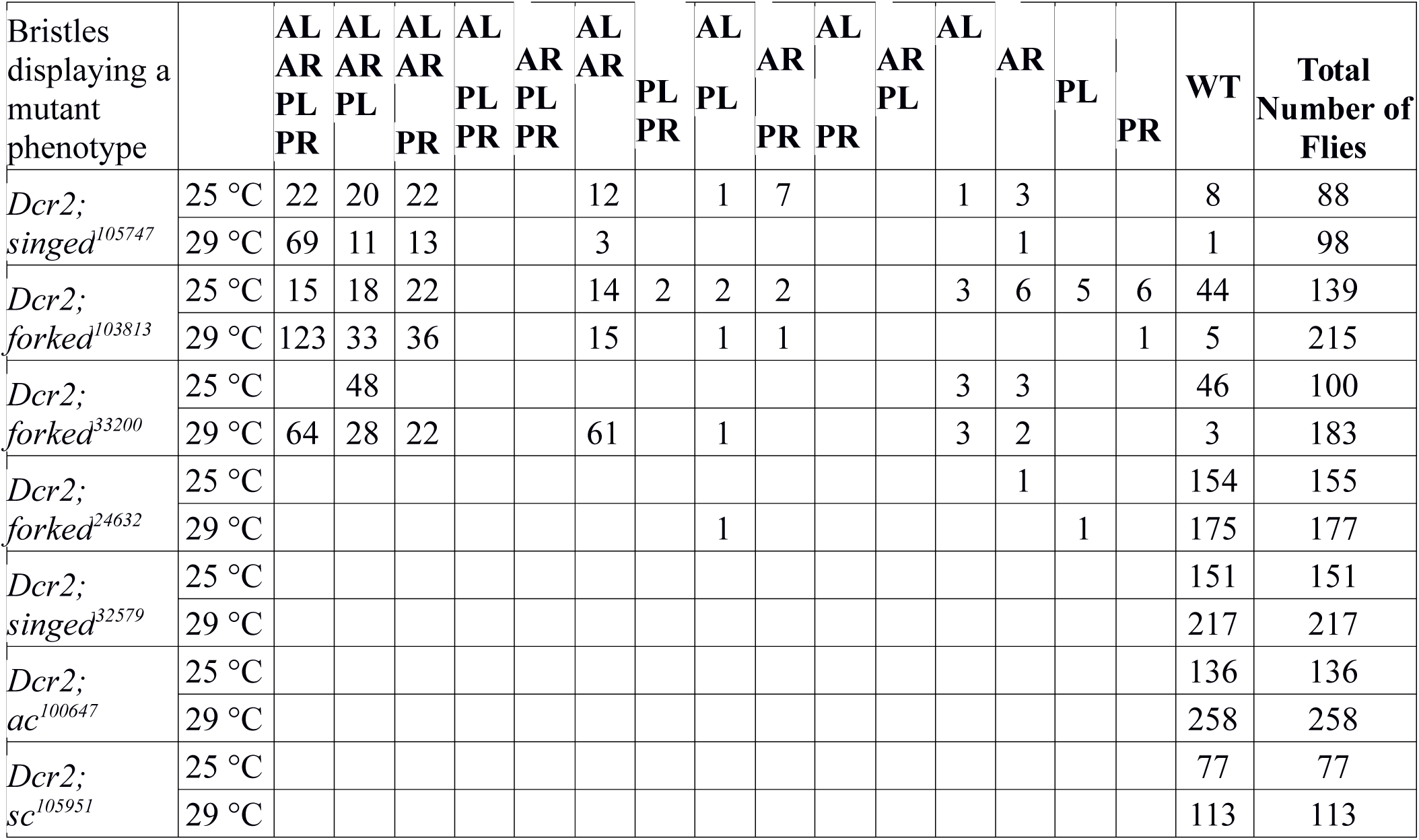
Test of various *UAS*-reporter constructs with *DC-GAL4*. The number of adults (males and females) with *singed/forked*/absent dorsocentral thoracic bristles is shown for various *UAS-RNAi* reporter lines. The phenotype was recorded individually for each of the four dorsocentral bristles: AL: anterior left, AR: anterior right, PL: posterior left, PR: posterior right. *Dcr2: UAS-Dicer-2*, WT: wild-type bristle phenotype. Crosses were performed at 25°C and at 29°C for each genotype.

**Table S6.**
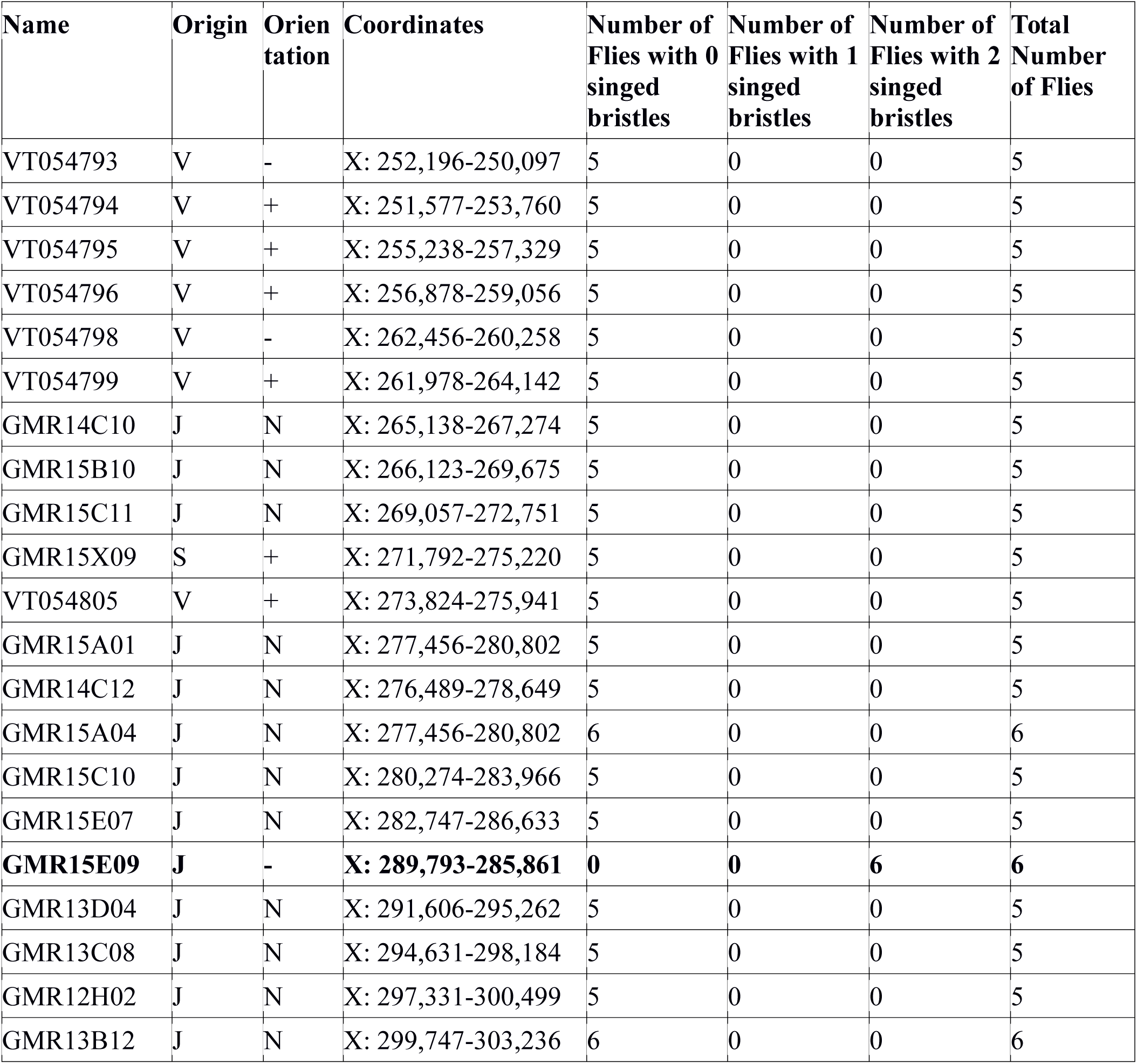
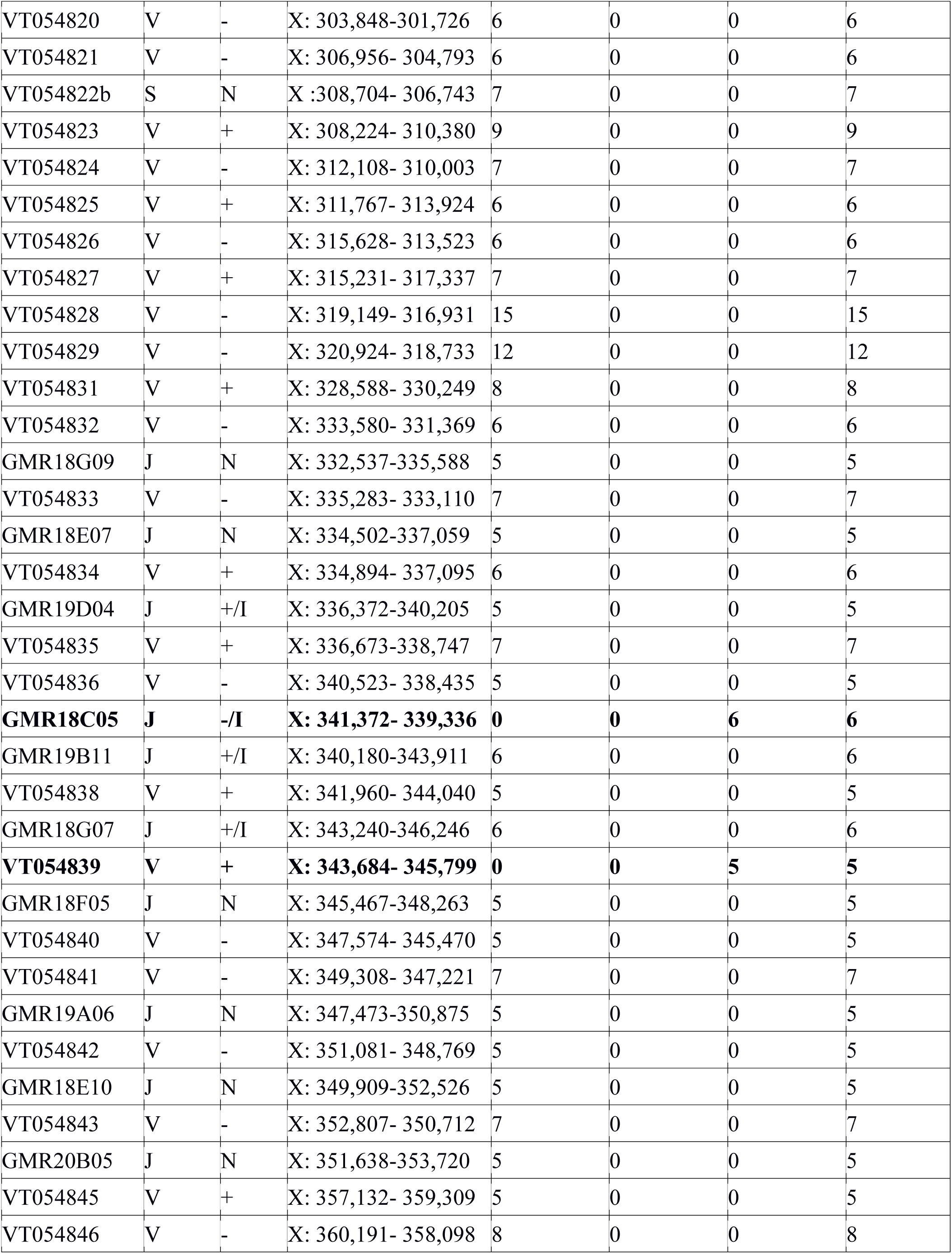
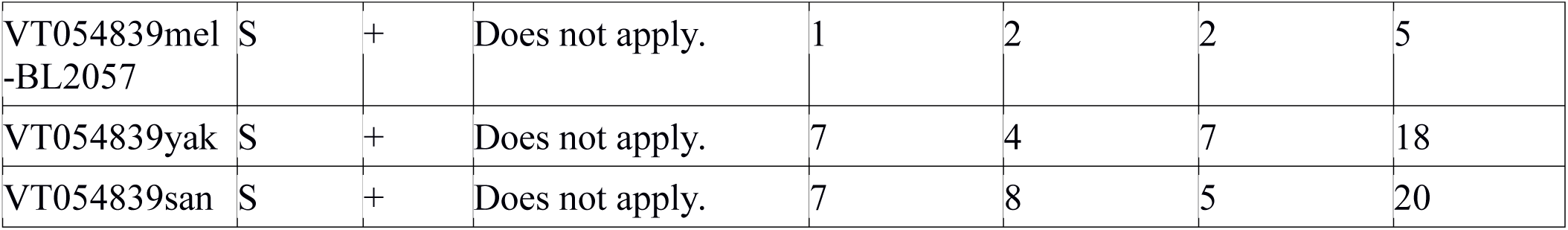
*Achaete-scute GAL4* lines and their hypandrial bristle phenotype with *Dcr2; UAS-singed.RNAi^105747^*. Lines are ordered according to their left coordinates (Release 5). The origin of each *GAL4* line is indicated by a letter code: V: Vienna Drosophila Research Center [63], J: Janelia Farm [38], S: This Study. The orientation of the *scute* cis-regulatory region within the reporter construct is indicated by + or -. N: no data, I: determined by PCR in this study. In bold are the 3 *GAL4* lines which produce a hypandrial bristle mutant phenotype. For short, *GMR15E09, GMR18C05* and *VT054839* are named *15E09, 18C05* and *054839* in the main text, respectively. Comparison of the *VT085439* cis-regulatory element from *D. santomea* and *D. yakuba* revealed no difference when tested in *GAL4* reporter lines with *Dcr2; UAS-singed.RNAi^105747^* (last two columns).

**Table S7.**
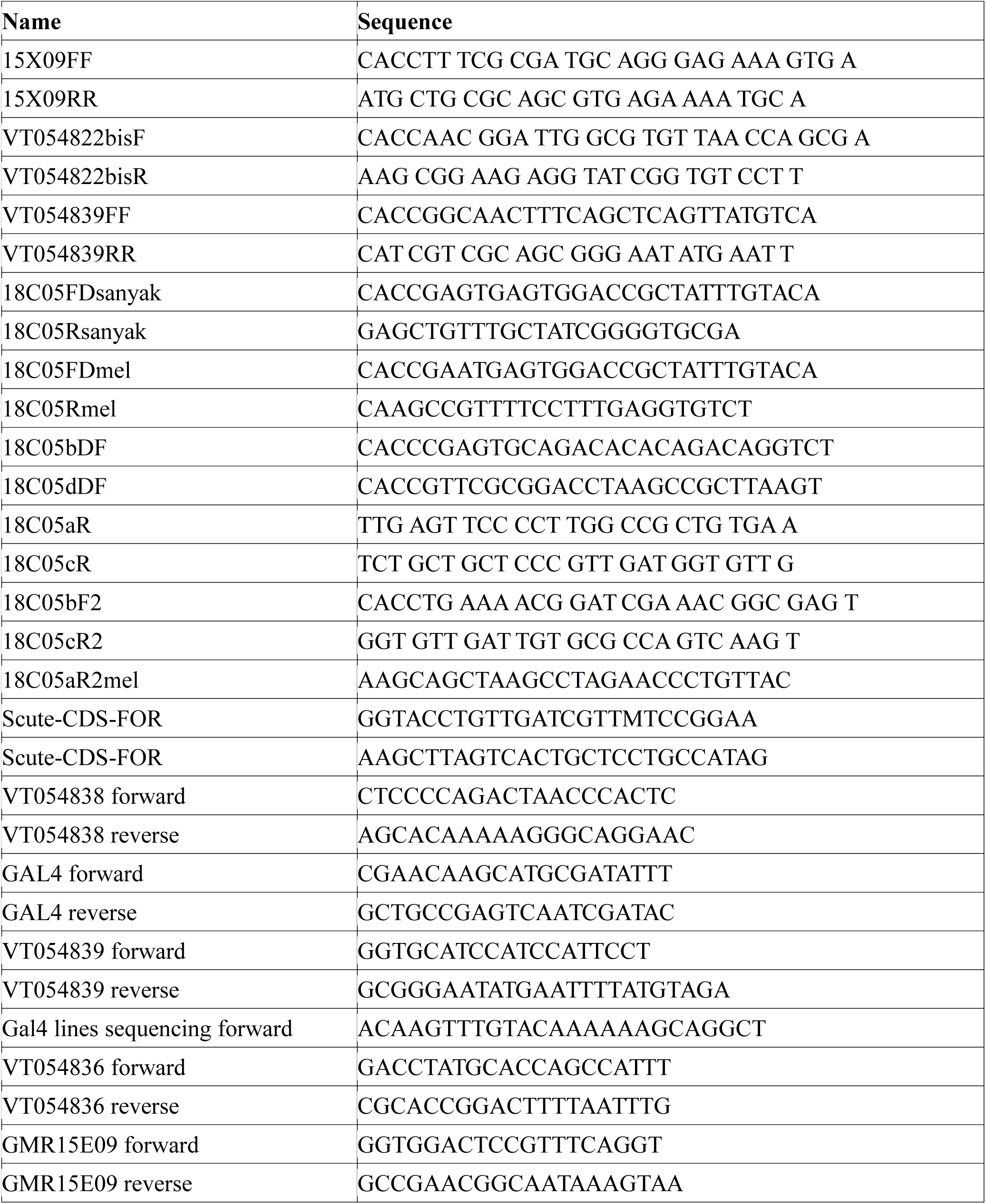
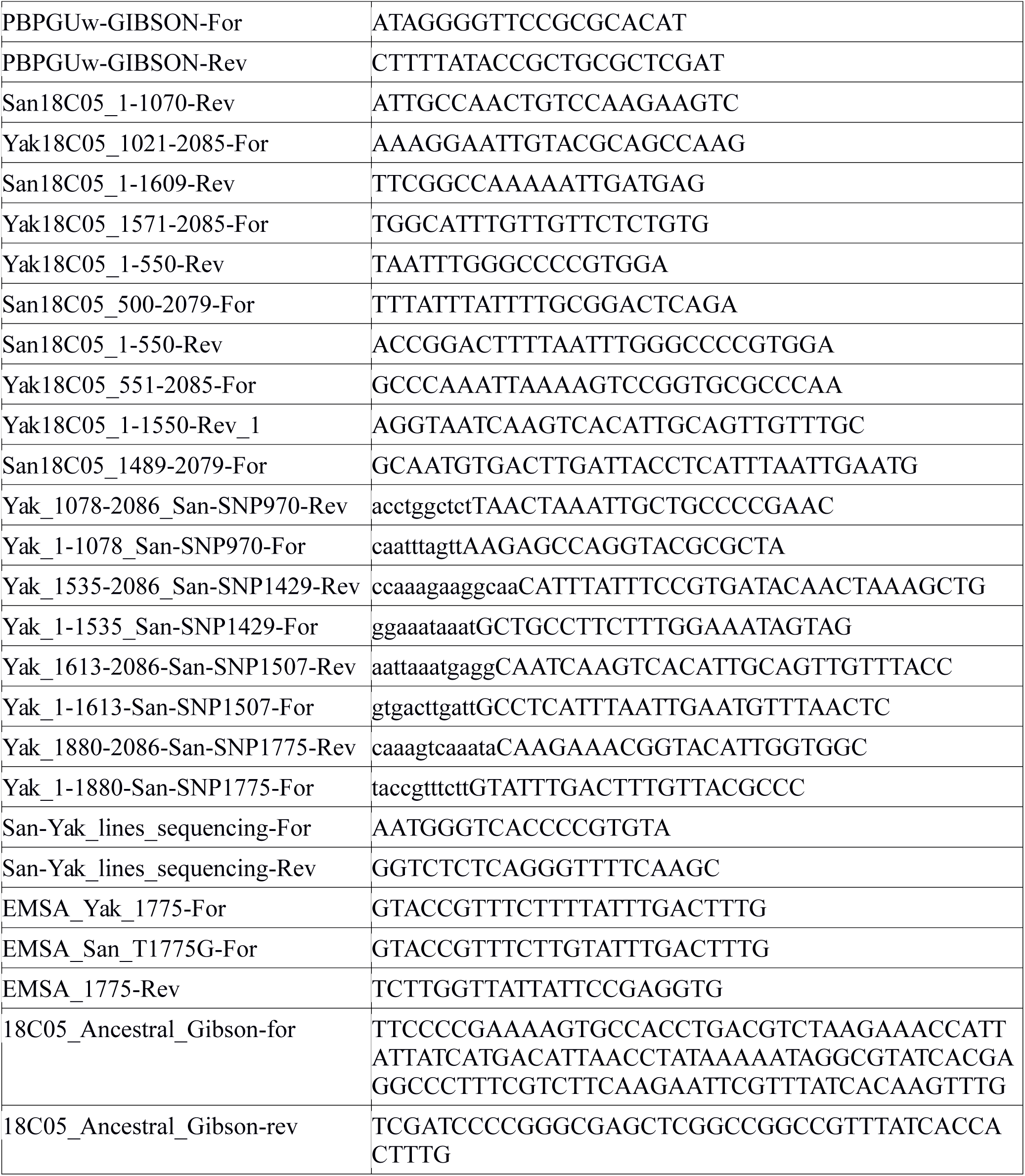
Primers used. All primers were purchased from Sigma Aldrich.

**Table S8.** PCR fragments cloned into pENTR/D-TOPO

| Fragment | Template | Forward primer | Reverse primer |
| --- | --- | --- | --- |
| 15X09 | *D. melanogaster* iso-1 | 15X09FF | 15X09RR |
| VT054822b | *D. melanogaster* iso-1 | VT054822bisF | VT054822bisR |
| VT054839yak | *D. yakuba* Ivory Coast | VT054839FF | VT054839RR |
| VT054839san | *D. santomea* SYN2005 | VT054839FF | VT054839RR |
| 18C05_T7 | *D. melanogaster* T7 | 18C05FDmel | 18C05Rmel |
| 18C05_BL2057 | *D. melanogaster* iso-1 | 18C05FDmel | 18C05Rmel |
| 18C05Yakfull | *D. yakuba* Ivory Coast | 18C05FDsanyak | 18C05Ryaksan |
| 18C05Sanfull | *D. santomea* SYN2005 | 18C05FDsanyak | 18C05Ryaksan |
| 18C05AmelBL | *D. melanogaster* iso-1 | 18C05FDmel | 18C05aR2mel |
| 18C05BmelBL | *D. melanogaster* iso-1 | 18C05bF2 | 18C05cR2 |
| 18C05CmelBL | *D. melanogaster* iso-1 | 18C05dDF | 18C05Rmel |
| 18C05ABmelBL | *D. melanogaster* iso-1 | 18C05FDmel | 18C05cR |
| 18C05BCmelBL | *D. melanogaster* iso-1 | 18C05bDF | 18C05Rmel |
| 18C05Asan | *D. santomea* SYN2005 | 18C05FDsanyak | 18C05aR |
| 18C05Bsan | *D. santomea* SYN2005 | 18C05bDF | 18C05cR2 |
| 18C05Ayak | *D. yakuba* Ivory Coast | 18C05FDsanyak | 18C05aR |
| 18C05Byak | *D. yakuba* Ivory Coast | 18C05bF2 | 18C05cR2 |
| 18C05Cyak | *D. yakuba* Ivory Coast | 18C05dDF | 18C05Ryaksan |
| VT054839mel-BL2057 | *D. melanogaster* iso-1 | VT054839FF | VT054839RR |
| VT054839yak | *D. yakuba* Ivory Coast | VT054839FF | VT054839RR |
| VT054839san | *D. santomea* SYN2005 | VT054839FF | VT054839RR |

**Table S9.**
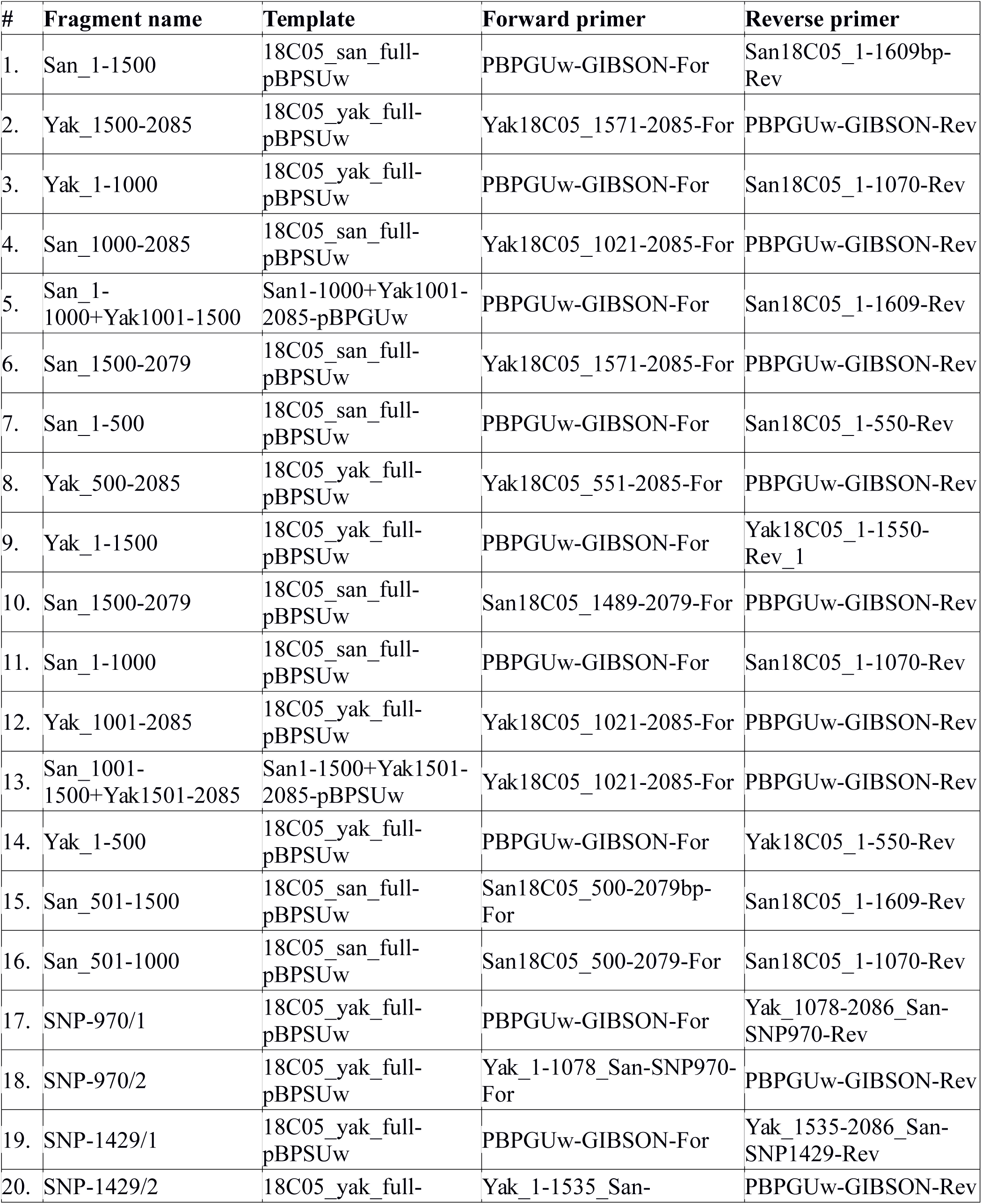
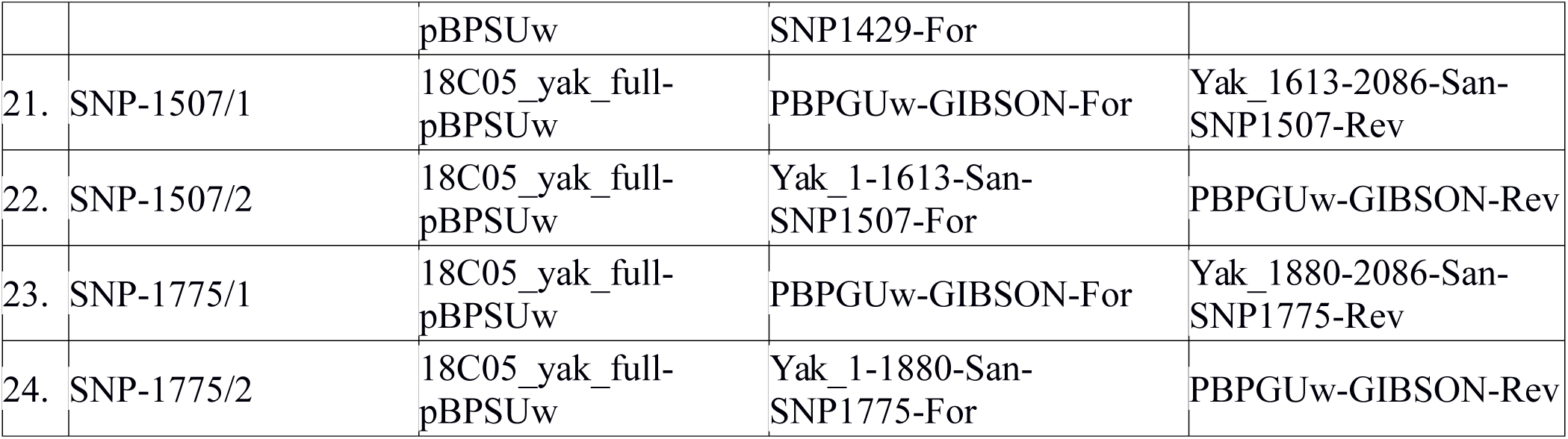
PCR fragments used for Gibson assembly into pBPSUw

**Table S10.** Scheme of Gibson assembly for chimeric constructs. See Table S9. for description of the PCR fragments.

| Construct Name | Assembled PCR Fragments |
| --- | --- |
| 18C05_SSSY | 1+2 |
| 18C05_YYSS | 3+4 |
| 18C05_SSYS | 5+6 |
| 18C05_SYYY | 7+8 |
| 18C05_YYYS | 9+10 |
| 18C05_SSYY | 11+12 |
| 18C05_YYSY | 3+13 |
| 18C05_YSYY | 14+16+12 |
| 18C05_YSSY | 14+15+2 |
| 18C05yakT970A | 17+18 |
| 18C05YyakT1429G | 19+20 |
| 18C05yakA1507G | 21+22 |
| 18C05yakt1775G | 23+24 |

**Table S11.**
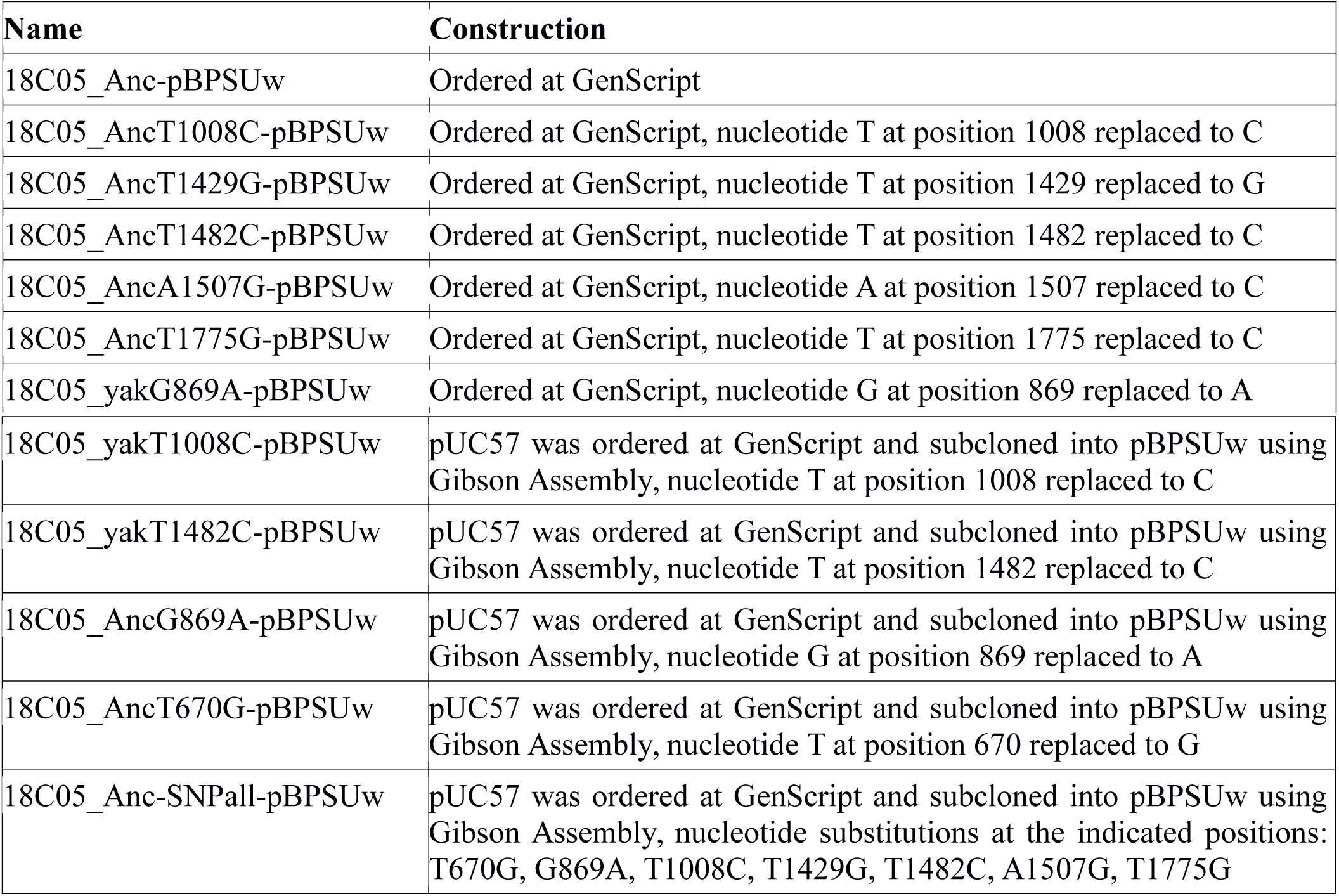
List of *18C05yakuba-* and *18C05ancestral-SNP-pBPSUw* constructs synthesized by GenScript

| Name | Construction |
| --- | --- |
| 18C05_Anc-pBPSUw | Ordered at GenScript |
| 18C05_AncT1008C-pBPSUw | Ordered at GenScript, nucleotide T at position 1008 replaced to C |
| 18C05_AncT1429G-pBPSUw | Ordered at GenScript, nucleotide T at position 1429 replaced to G |
| 18C05_AncT1482C-pBPSUw | Ordered at GenScript, nucleotide T at position 1482 replaced to C |
| 18C05_AncA1507G-pBPSUw | Ordered at GenScript, nucleotide A at position 1507 replaced to C |
| 18C05_AncT1775G-pBPSUw | Ordered at GenScript, nucleotide T at position 1775 replaced to C |
| 18C05_yakG869A-pBPSUw | Ordered at GenScript, nucleotide G at position 869 replaced to A |
| 18C05_yakT1008C-pBPSUw | pUC57 was ordered at GenScript and subcloned into pBPSUw using Gibson Assembly, nucleotide T at position 1008 replaced to C |
| 18C05_yakT1482C-pBPSUw | pUC57 was ordered at GenScript and subcloned into pBPSUw using Gibson Assembly, nucleotide T at position 1482 replaced to C |
| 18C05_AncG869A-pBPSUw | pUC57 was ordered at GenScript and subcloned into pBPSUw using Gibson Assembly, nucleotide G at position 869 replaced to A |
| 18C05_AncT670G-pBPSUw | pUC57 was ordered at GenScript and subcloned into pBPSUw using Gibson Assembly, nucleotide T at position 670 replaced to G |
| 18C05_Anc-SNPall-pBPSUw | pUC57 was ordered at GenScript and subcloned into pBPSUw using Gibson Assembly, nucleotide substitutions at the indicated positions: T670G, G869A, T1008C, T1429G, T1482C, A1507G, T1775G |

**Table S12.**
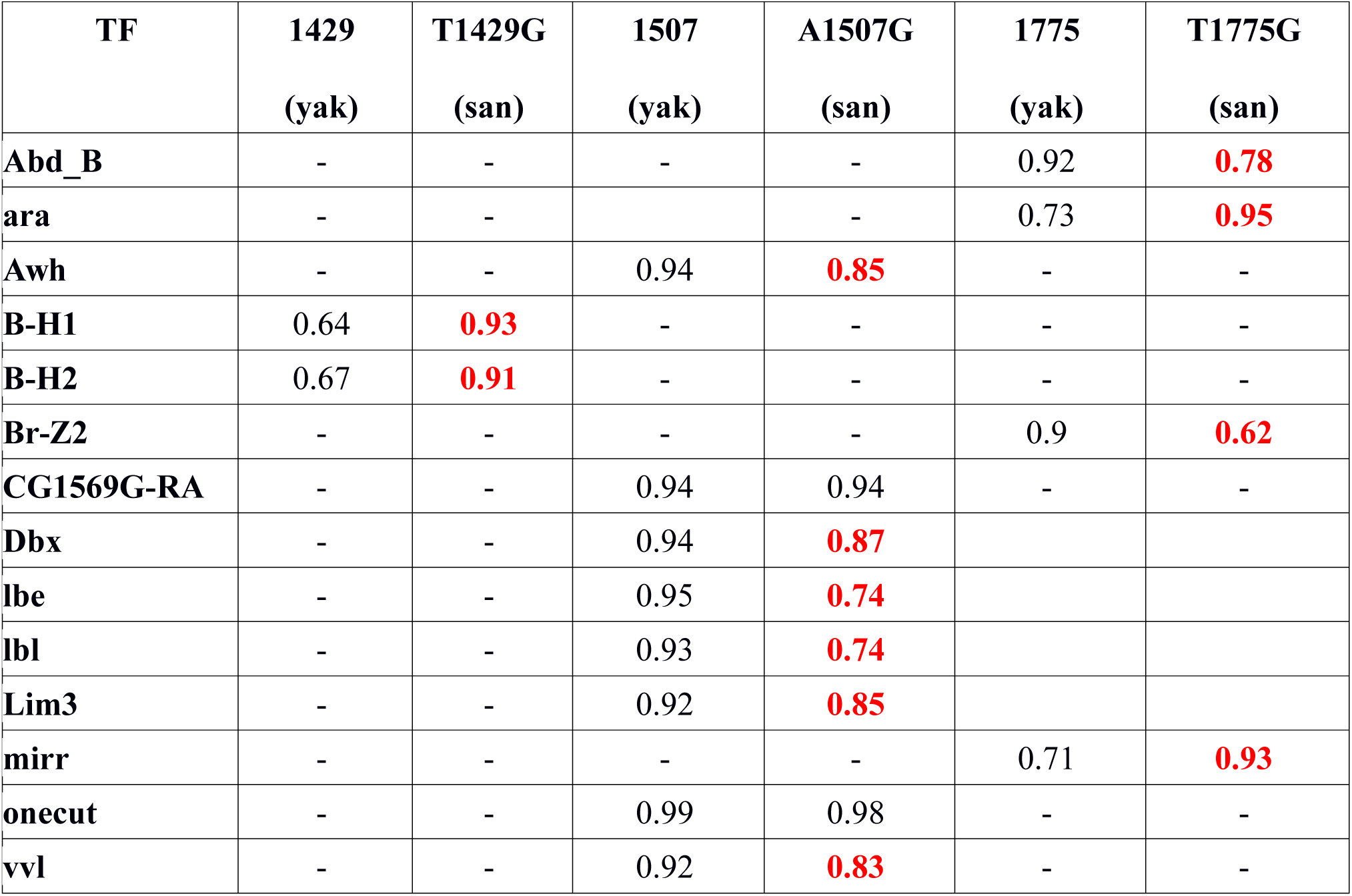
JASPAR binding scores for 6 loci of interest within the *18C05* region. Predicted transcription factors (TF) binding sites and their binding scores are shown for 3 loci of interest within the *18C05* region which underwent *D. santomea-specific* substitutions associated with a decrease in hypandrial bristle number For each position the left column indicates binding scores for the *D. yakuba* sequence and the right column for the *D. santomea* sequence. All the TF predicted by JASPAR with binding scores higher than 0.9 for at least one locus are shown. “-” indicates cases where binding scores of the *D. yakuba* sequence is less than 0.9. In red are binding scores that were changed by the mutation. We found for T1429G a gain of two repressors (B-H1 and B-H2), for A1507G a loss of several TF binding sites (AWH, DDX, LBE, LBL, LIM3 and VVL), for T1775G a gain of ARA and MIRR binding and a loss of BR-Z and ABD-B binding sites.

**Table S13.** Hypandrial bristle phenotypes in *ABD-B RNAi* lines and mitotic clones. *Abd-B RNAi* was induced with different *GAL4* drivers and *Abd-B^M1^* and *Abd-B^D18^* mutations were used in mitotic clones. The number of dissected flies and recognizable hypandrium with a shape similar to wild-type are shown. The number of hypandrium with 0, 1, 2, 3 or with 2 misplaced bristles are shown in separate columns. Different bristle phenotypes are indicated with the following labels: n: normal, t: thin, s: shorter than wild-type, M: Minute (thinner and slightly shorter than wild-type), y+: non yellow, y-: yellow, R: 2 bristles on the right, L: 2 bristles on the left.

| Genotype | Number of dissected flies | Number of recognizable hypandrium | 0 bristle | 1 bristle | 2 bristles | 3 bristles | 2 misplaced bristle |
| --- | --- | --- | --- | --- | --- | --- | --- |
| *Abd-B.RNAi<sup>26746</sup>/NP6333-GAL4* | 50 | 0 |  |  |  |  |  |
| *Abd-B.RNAi<sup>26746</sup>/NP5130-GAL4* | 50 | 0 |  |  |  |  |  |
| *Abd-B.RNAi<sup>51167</sup>/NP6333-GAL4* | 50 | 9 |  | 1(t,s) | 6(t,s) | 1(t,s) | 1(L) |
| *Abd-B.RNAi<sup>51167</sup>/NP5130-GAL4* | 50 | 11 |  |  | 9(t,s) |  | 1(L), 1(R) |
| *Abd-B.RNAi<sup>51167</sup>/GMR18C05-GAL4* | 10 | 10 |  |  | 7(n), 3(t) |  |  |
| *yw hsflp122; FRT82B hs-CD2 y+ M(3) w<sup>123</sup> /FRT82B Abd-B<sup>M1</sup> red[1] e[11] ro[1] ca[1]* | 30 | 10 | 2 | 2(M,y<sup>+</sup>) | 2(M,y<sup>+</sup>),<br>1(M,y<sup>-</sup>),<br>1(M,y<sup>+</sup>; M,y<sup>-</sup>) |  | 2(R,M,y<sup>-</sup>) |
| *yw hsflp122; FRT82B hs-CD2 y+ M(3) w<sup>123</sup> /FRT82B Abd-B<sup>D18</sup>* | 52 | 2 | 1 | 1(y<sup>+</sup>) |  |  |  |

## AUXILIARY DATA FILES

**Data S1. QTL-data (separate file)**

Data for Fig. 2.

**Data S2. Hypandrial bristle number with *18C05-GAL4* constructs (separate file)**

Data for Fig. 3.

**Data S3. Hypandrial bristle number with *18C05-sc* constructs (separate file)**

Data for Fig. 3., Fig S3., Fig S4.

**Data S4. Sex comb tooth number with various *18C05yak-sc* constructs (separate file)**

Data for Fig. 4.

**Data S5. Genital bristle numbers in *D. yakuba* and *D. santomea* strains (separate file)**

Data for Fig. S1.

**Data S6. Hypandrial bristle number with *18C05 melanogaster-GAL4* constructs (separate file)**

Data for Fig. S5.

**Data S7. Hypandrial bristle number with chimeric constructs (separate file)**

Data for Fig. S8.

**Data S8. Sex comb tooth numbers in *D. yakuba, D. santomea* and in *D. melanogaster* Canton S and sc mutants (separate file)**

Data for Fig. S9.

**Data S9. Fractional occupancy data (separate file)**

Data for Fig. S11. EMSA shift intensity values measured by ImageJ and the calculated fractional occupancy.

**Data S10. Number of GFP-positive cells in 5h APF pupal legs of *D. melanogaster* flies of genotype *18C05yak-GFP* or *18C05yakT1775G-GFP* or *18C05san-GFP* (separate file)**

Data for Fig. 4.

## REFERENCES

1. Darwin, C. (1859). On the origin of species by means of natural selection, or the preservation of favoured races in the struggle for life.

2. Saltz, J.B., Hessel, F.C., and Kelly, M.W. (2017). Trait Correlations in the Genomics Era. Trends Ecol. Evol. (Amst.) 32, 279–290.

3. Paaby, A.B., and Rockman, M.V. (2013). The many faces of pleiotropy. Trends in Genetics 29, 66–73.

4. Fisher, R.A. (1930). The genetical theory of natural selection (Oxford: Clarendon).

5. Orr, H.A. (2000). Adaptation and the cost of complexity. Evolution 54, 13–20.

6. Wagner, G.P., and Zhang, J. (2011). The pleiotropic structure of the genotype–phenotype map: the evolvability of complex organisms. Nature Reviews Genetics 12, 204–213.

7. Stearns, F.W. (2010). One Hundred Years of Pleiotropy: A Retrospective. Genetics 186, 767–773.

8. Lonfat, N., Montavon, T., Darbellay, F., Gitto, S., and Duboule, D. (2014). Convergent evolution of complex regulatory landscapes and pleiotropy at Hox loci. Science 346, 1004–1006.

9. Preger-Ben Noon, E., Sabarís, G., Ortiz, D.M., Sager, J., Liebowitz, A., Stern, D.L., and Frankel, N. (2018). Comprehensive Analysis of a cis -Regulatory Region Reveals Pleiotropy in Enhancer Function. Cell Reports 22, 3021–3031.

10. Duveau, F., and Félix, M.-A. (2012). Role of pleiotropy in the evolution of a cryptic developmental variation in Caenorhabditis elegans. PLoS Biol. 10, e1001230.

11. Chang, S.H., Jobling, S., Brennan, K., and Headon, D.J. (2009). Enhanced Edar Signalling Has Pleiotropic Effects on Craniofacial and Cutaneous Glands. PLOS ONE 4, e7591.

12. Kent, C.F., Daskalchuk, T., Cook, L., Sokolowski, M.B., and Greenspan, R.J. (2009). The Drosophila foraging Gene Mediates Adult Plasticity and Gene–Environment Interactions in Behaviour, Metabolites, and Gene Expression in Response to Food Deprivation. PLOS Genetics 5, e1000609.

12b. Endler, L., Gibert, J. M., Nolte, V., & Schlötterer, C. (2018). Pleiotropic effects of regulatory variation in tan result in correlation of two pigmentation traits in *Drosophila melanogaster*. Molecular ecology

13. Wittkopp, P.J., Haerum, B.K., and Clark, A.G. (2008). Regulatory changes underlying expression differences within and between Drosophila species. Nature Genetics 40, 346–350.

14. Turissini, D.A., and Matute, D.R. (2017). Fine scale mapping of genomic introgressions within the Drosophila yakuba clade. PLoS Genetics 13, e1006971.

15. Lachaise, D., Harry, M., Solignac, M., Lemeunier, F., Bénassi, V., and Cariou, M.L. (2000). Evolutionary novelties in islands: Drosophila santomea, a new melanogaster sister species from São Tomé. Proc. Biol. Sci. 267, 1487–1495.

16. Simpson, P., Woehl, R., and Usui, K. (1999). The development and evolution of bristle patterns in Diptera. Development 126, 1349–1364.

17. Gómez-Skarmeta, J.L., Rodríguez, I., Martínez, C., Culí, J., Ferrés-Marcó, D., Beamonte, D., and Modolell, J. (1995). Cis-regulation of achaete and scute: shared enhancer-like elements drive their coexpression in proneural clusters of the imaginal discs. Genes Dev 9, 1869–1882.

18. Marcellini, S., Gibert, J.-M., and Simpson, P. (2005). achaete, but not scute, is dispensable for the peripheral nervous system of Drosophila. Dev. Biol. 285, 545–553.

19. Jory, A., Estella, C., Giorgianni, M.W., Slattery, M., Laverty, T.R., Rubin, G.M., and Mann, R.S. (2012). A Survey of 6,300 Genomic Fragments for cis-Regulatory Activity in the Imaginal Discs of Drosophila melanogaster. Cell Reports 2, 1014–1024.

19b. Tanaka, K., Barmina, O., Sanders, L. E., Arbeitman, M. N., & Kopp, A. (2011). Evolution of sex-specific traits through changes in HOX-dependent doublesex expression. PLoS biology, 9(8), e1001131.

20. Ng, C.S., and Kopp, A. (2008). Sex combs are important for male mating success in Drosophila melanogaster. Behavior genetics 38, 195.

21. Coyne, J.A., Elwyn, S., Kim, S.Y., and Llopart, A. (2004). Genetic studies of two sister species in the Drosophila melanogaster subgroup, D. yakuba and D. santomea. Genetics Research 84, 11–26.

22. Foronda, D., Estrada, B., de Navas, L., and Sánchez-Herrero, E. (2006). Requirement of Abdominal-A and Abdominal-B in the developing genitalia of Drosophila breaks the posterior downregulation rule. Development 133, 117–127.

23. Hurtado-Gonzales, J.L., Gallaher, W., Warner, A., and Polak, M. (2015). Microscale Laser Surgery Demonstrates the Grasping Function of the Male Sex Combs in Drosophila melanogaster and Drosophila bipectinata. Ethology 121, 45–56.

24. Eberhard, W.G. (1988). Sexual Selection and Animal Genitalia (Harvard University Press).

25. McLean, C.Y., Reno, P.L., Pollen, A.A., Bassan, A.I., Capellini, T.D., Guenther, C., Indjeian, V.B., Lim, X., Menke, D.B., Schaar, B.T., et al. (2011). Human-specific loss of regulatory DNA and the evolution of human-specific traits. Nature 471, 216–219.

26. Mayr, E. (1963). Animal species and evolution (Harvard University Press).

27. Carroll, S.B. (2008). Evo-devo and an expanding evolutionary synthesis: a genetic theory of morphological evolution. Cell 134, 25–36.

28. Cheng, Y., Ma, Z., Kim, B.-H., Wu, W., Cayting, P., Boyle, A.P., Sundaram, V., Xing, X., Dogan, N., Li, J., et al. (2014). Principles of regulatory information conservation between mouse and human. Nature 515, 371.

## SUPPLEMENTAL REFERENCES

29. Andolfatto, P., Davison, D., Erezyilmaz, D., Hu, T.T., Mast, J., Sunayama-Morita, T., and Stern, D.L. (2011). Multiplexed shotgun genotyping for rapid and efficient genetic mapping. Genome Res. 21, 610–617.

30. Broman, K.W., and Sen, S. (2009). A Guide to QTL Mapping with R/qtl 1st ed. (Springer).

31. Broman, K.W., Wu, H., Sen, S., and Churchill, G.A. (2003). R/qtl: QTL mapping in experimental crosses. Bioinformatics 19, 889–890.

32. Haley, C.S., and Knott, S.A. (1992). A simple regression method for mapping quantitative trait loci in line crosses using flanking markers. Heredity (Edinb) 69, 315–324.

33. Venken, K.J.T., Popodi, E., Holtzman, S.L., Schulze, K.L., Park, S., Carlson, J.W., Hoskins, R.A., Bellen, H.J., and Kaufman, T.C. (2010). A molecularly defined duplication set for the X chromosome of Drosophila melanogaster. Genetics 186, 1111–1125.

34. Turissini, D.A., McGirr, J.A., Patel, S.S., David, J.R., and Matute, D.R. (2017). The rate of evolution of postmating-prezygotic reproductive isolation in Drosophila. Mol. Biol. Evol.

35. Schneider, C.A., Rasband, W.S., and Eliceiri, K.W. (2012). NIH Image to ImageJ: 25 years of image analysis. Nature methods 9, 671–675.

36. Taylor, B.J. (1989). Sexually dimorphic neurons in the terminalia of Drosophila melanogaster: I. Development of sensory neurons in the genital disc during metamorphosis. J. Neurogenet. 5, 173–192.

37. Dietzl, G., Chen, D., Schnorrer, F., Su, K.-C., Barinova, Y., Fellner, M., Gasser, B., Kinsey, K., Oppel, S., Scheiblauer, S., et al. (2007). A genome-wide transgenic RNAi library for conditional gene inactivation in Drosophila. Nature 448, 151–156.

38. Jenett, A., Rubin, G.M., Ngo, T.-T., Shepherd, D., Murphy, C., Dionne, H., Pfeiffer, B.D., Cavallaro, A., Hall, D., Jeter, J., et al. (2012). A GAL4-driver line resource for Drosophila neurobiology. Cell reports 2, 991–1001.

39. Gibson, D.G., Young, L., Chuang, R.-Y., Venter, J.C., Hutchison, C.A., and Smith, H.O. (2009). Enzymatic assembly of DNA molecules up to several hundred kilobases. Nature methods 6, 343–345.

40. Bryne, J.C., Valen, E., Tang, M.-H.E., Marstrand, T., Winther, O., da Piedade, I., Krogh, A., Lenhard, B., and Sandelin, A. (2007). JASPAR, the open access database of transcription factor-binding profiles: new content and tools in the 2008 update. Nucleic acids research 36, D102–D106.

41. Zhu, L.J., Christensen, R.G., Kazemian, M., Hull, C.J., Enuameh, M.S., Basciotta, M.D., Brasefield, J.A., Zhu, C., Asriyan, Y., Lapointe, D.S., et al. (2010). FlyFactorSurvey: a database of Drosophila transcription factor binding specificities determined using the bacterial one-hybrid system. Nucleic acids research 39, D111–D117.

42. Crawley, M.J. (2012). The R book (John Wiley & Sons).

43. Hilbe, J.M. (2014). Modeling Count Data (Cambridge University Press).

44. Zuur, A., Ieno, E.N., Walker, N., Saveliev, A.A., and Smith, G.M. (2011). Mixed Effects Models and Extensions in Ecology with R Softcover reprint of hardcover 1st ed. 2009 edition. (New York, NY: Springer).

45. Team, R.C. (2016). R: A language and environment for statistical (Vienna, Austria) Available at: http://www.R-project.org/.

46. Bates, D., Mächler, M., Bolker, B., and Walker, S. (2014). Fitting linear mixed-effects models using lme4. arXiv preprint arXiv:1406.5823.

47. Hothorn, T., Bretz, F., and Westfall, P. (2008). Simultaneous inference in general parametric models. Biometrical journal 50, 346–363.

48. Bretz, F., Hothorn, T., and Westfall, P. (2010). Multiple comparisons using R (CRC Press).

49. Holm, S. (1979). A simple sequentially rejective multiple test procedure. Scandinavian journal of statistics, 65–70.

50. Frangioni, J.V., and Neel, B.G. (1993). Solubilization and purification of enzymatically active glutathione S-transferase (pGEX) fusion proteins. Analytical biochemistry 210, 179–187.

51. Fan, Y.-J., Gittis, A.H., Juge, F., Qiu, C., Xu, Y.-Z., and Rabinow, L. (2014). Multifunctional RNA Processing Protein SRm160 Induces Apoptosis and Regulates Eye and Genital Development in Drosophila. Genetics 197, 1251–1265.

52. Chatterjee, S.S., Uppendahl, L.D., Chowdhury, M.A., Ip, P.-L., and Siegal, M.L. (2011). The female-specific doublesex isoform regulates pleiotropic transcription factors to pattern genital development in Drosophila. Development 138, 1099–1109.

53. Skaer, N., Pistillo, D., and Simpson, P. (2002). Transcriptional heterochrony of scute and changes in bristle pattern between two closely related species of blowfly. Developmental biology 252, 31–45.

54. Casanova, J., Sánchez-Herrero, E., and Morata, G. (1986). Identification and characterization of a parasegment specific regulatory element of the abdominal-B gene of Drosophila. Cell 47, 627–636.

55. Hopmann, R., Duncan, D., and Duncan, I. (1995). Transvection in the iab-5, 6, 7 region of the bithorax complex of Drosophila: homology independent interactions in trans. Genetics 139, 815–833.

56. Xu, T., and Rubin, G.M. (1993). Analysis of genetic mosaics in developing and adult Drosophila tissues. Development 117, 1223–1237.

57. Estrada, B., and Sánchez-Herrero, E. (2001). The Hox gene Abdominal-B antagonizes appendage development in the genital disc of Drosophila. Development 128, 331–339.

58. Maroni, G. and S.C. Stamey (1983). Developmental profile and tissue distribution of alcohol dehydrogenase. Drosophila Information Service 59.

59. Andres, A.J., and Thummel, C.S. (1994). Methods for quantitative analysis of transcription in larvae and prepupae. Methods Cell Biol. 44, 565–573.

59b. Corson, F., Couturier, L., Rouault, H., Mazouni, K., & Schweisguth, F. (2017). Self-organized Notch dynamics generate stereotyped sensory organ patterns in Drosophila. Science, 356(6337), eaai7407.

60. Campuzano, S., Carramolino, L., Cabrera, C.V., Ruíz-Gómez, M., Villares, R., Boronat, A., and Modolell, J. (1985). Molecular genetics of the achaete-scute gene complex of D. melanogaster. Cell 40, 327–338.

61. Carramolino, L., Ruiz-Gomez, M., del Carmen Guerrero, M., Campuzano, S., and Modolell, J. (1982). DNA map of mutations at the scute locus of Drosophila melanogaster. The EMBO journal 1, 1185.

62. Cubas, P., De Celis, J.F., Campuzano, S., and Modolell, J. (1991). Proneural clusters of achaete-scute expression and the generation of sensory organs in the Drosophila imaginal wing disc. Genes & Development 5, 996–1008.

63. Kvon, E.Z., Kazmar, T., Stampfel, G., Yáñez-Cuna, J.O., Pagani, M., Schernhuber, K., Dickson, B.J., and Stark, A. (2014). Genome-scale functional characterization of Drosophila developmental enhancers in vivo. Nature 512, 91–95., 91-95.

